# Genetic tracing of the white-bellied pangolin’s trade in western central Africa

**DOI:** 10.1101/2023.03.10.530129

**Authors:** Alain Din Dipita, Alain Didier Missoup, Samantha Aguillon, Emilie Lecompte, Brice Roxan Momboua, Anne-Lise Chaber, Katharine Abernethy, Flobert Njiokou, Maurice Tindo, Stephan Ntie, Philippe Gaubert

## Abstract

African pangolins are intensively harvested to feed illegal trade networks. We focused on the conservation genetics of the most trafficked African species, the white-bellied pangolin (WBP; *Phataginus tricuspis*), through the genotyping of 562 individuals from reference populations and urban bushmeat markets in a vibrant trade hub from western Central Africa. Across Cameroon, Equatorial Guinea and northern Gabon, we observed a lack of genetic differentiation and a signature of isolation-by-distance possibly due to unsuspected dispersal capacities involving a Wahlund effect. Despite a higher level of genetic diversity compared to western Africa, we detected a 74-83% decline in the effective population size of WBP during the Middle Holocene. Private allele frequency tracing approach indicated up to 600 km sourcing distance by large urban markets from Cameroon, involving transnational trade activities. The 20 microsatellites markers used in this study provided the necessary power to distinguish among all WBP individuals and should be considered a valuable resource for future forensic applications. Because lineage admixture was detected in the study area, we recommend a multi- locus approach for tracing the WBP trade. The Yaoundé market was a major recruiter of genetic diversity in the region, and should receive urgent conservation action to mitigate the pangolin trade.

## Introduction

The illegal wildlife trade is a flourishing and parallel economy, representing a global annual revenue of US$7-23 billion (Morton et al., 2021; Sas-Rolfes et al., 2019). It is fed by uncontrolled hunting activities mostly affecting the tropics, with a significant and deleterious impact on the survival and abundance of terrestrial vertebrates (Benítez-López et al., 2017; Maxwell et al., 2016). Pangolins (Mammalia, Pholidota) have been spotlighted as one of the emblems of over-harvesting due to the illegal wildlife trade, being considered the most trafficked mammals in the world (Challender et al., 2015, 2020; Heinrich et al., 2017). With approximately 900,000 individuals seized in the last 20 years (Challender et al., 2020), the eight species of African and Asian pangolins are threatened with extinction, through the cumulative effect of illegal hunting and deforestation (Heighton & Gaubert, 2021). As a consequence, they have all been listed on Appendix I of the Convention on International Trade in Endangered Species of Wild Fauna and Flora (CITES) (Challender & Watermann, 2017), and rated as Vulnerable, Endangered or Critically Endangered on the IUCN Red List of Threatened Species™ (Hoffmann & Challender, 2020).

In Africa, pangolins have long been hunted as part of the traditional bushmeat spectrum used by local communities (Boakye et al., 2016; Zanvo et al., 2021). However, the apparent decrease in numbers of Asian pangolins (Challender & Watermann, 2017; Heinrich et al., 2017) and the large demand from the Chinese traditional medicine market (Challender & Watermann, 2017; Cheng et al., 2017), have recently set up a global trade network where African pangolins are being massively exported –mostly for their scales– to South-East Asia (Challender et al., 2012; Ingram et al., 2019; Zhang et al., 2020). Challender et al. (2020) estimated that > 400,000 African pangolins were bound for Asian markets between 2015 and 2019. In Africa, a 42yr- records survey suggests that harvest volumes of pangolins were multiplied by nine between 2005 and 2014 (Ingram & Coad, 2016). For central Africa alone, the amount of yearly extirpated pangolins has been estimated to increase from 0.42 to 2.71 M (Ingram et al., 2018). Despite pangolins having been –erroneously– blamed for their role as potential intermediate hosts of the COVID-19 pandemic (Frutos et al., 2020), African pangolins continue to be harvested at high rates (Aditya et al., 2021).

Most of the efforts to trace the pangolin trade has been focusing on species identification, using different types of samples and DNA quality (see Kotze et al., 2020). The genetic tracing of the pangolin trade has proven efficient in species identification (Ewart et al., 2021; Gaubert & Antunes, 2015; Singh et al., 2021) but has remained limited in reaching lineage or population-level assignment, notably because of incomplete DNA registers (Baker, 2008). Genetic resources recently produced for the white bellied pangolin (WBP; *Phataginus tricuspis*), the commonest and most traded species in Africa (Heinrich et al., 2016), have provided a unique opportunity to trace the pangolin trade at both the local and global scales. Using mitochondrial and nuclear DNA sequences, Gaubert et al. (2016) delineated six cryptic, divergent and geographically isolated lineages across the species’ range, that were later traced by Zhang et al. (2020) from large scale seizures made in Hong Kong. Further development of specific microsatellites markers (Aguillon et al., 2020) and extensive sample collection allowed to trace the WBP trade at the scale of a sub-region, the Dahomey Gap (West Africa), demonstrating 200–300 km trade distances between sourced forest and urban markets (Zanvo et al., 2022).

Despite the original development of microsatellite markers from Cameroonian samples (Aguillon et al., 2020), no extensive genetic studies have yet been conducted on pangolins in the country. Cameroon has a long history of bushmeat consumption and trade (Bahuchet & Ioveva, 1999), and pangolins are commonly offered for sale in bushmeat markets and restaurants (Ingram et al., 2018; Nguyen et al., 2021), even since the advent of the COVID-19 pandemic (Harvey-Carroll et al., 2022). Although pangolins have been nationally protected since 2016, Cameroon remains one of the hotspots of the illegal pangolin trade, exporting a large amount of scales from Africa to Asia (Ingram et al., 2019), with WBP being by far the main contributing species (Pietersen & Challender, 2019).

Cameroon is home to the Western Central Africa (WCA) lineage (Gaubert et al., 2016), also extending southward to Equatorial Guinea and northern Gabon, and which was the most represented lineage in recent Asian seizures (Zhang et al., 2020). Given the uncontrolled domestic and international trade affecting the species, tracing the trade of WBP in Cameroon and neighboring countries is a stake with primordial conservation implications. WCA is co-occurring in Gabon with the Gab lineage, which was temporarily considered as endemic to the country pending more extensive geographic sampling (Gaubert et al., 2016). Because (i) admixture between pangolin lineages may jeopardize efforts at tracing the pangolin trade (Nash et al., 2018) and (ii) both WCA and Gab have been detected in seizures from Hong Kong (Zhang et al., 2020), we propose to build on recently developed genetic resources to investigate both maternal and bi-parental population genetic signatures in WBP from the subregion. Our specific objectives were to (i) assess the population structure and genetic diversity of WBP from Cameroon, Equatorial Guinea and Gabon, (ii) characterize their demographic history and current effective population size, and (iii) trace the scale of the sub-regional WBP trade.

## Methods

### Genetic sampling and wet laboratory procedures

We collected 565 genetic samples of WBP from the Western Central Africa lineage (WCA), across Cameroon, Equatorial Guinea (including Bioko Island) and Gabon between 2007 and 2019. Because the endemic Gabon lineage (Gab) co-occurs with WCA in Gabon (Gaubert et al., 2016), we assume that samples from Gab were also collected. We surveyed and collected 188 samples from 37 local bushmeat markets and forest sites, and 377 samples from three large urban bushmeat markets in Cameroon (two in Douala and one in Yaoundé) and from seized luggage originating from Cameroonian flights at Roissy (Paris, France) and Brussels (Belgium) airports (see Supplementary Fig. S1 and Supplementary Table S1 online). We collected through upstream interviews with urban market sellers’ information on the geographic origin of the pangolins sold. Using the same approach with local market vendors, we could confirm that all the pangolins collected on the local bushmeat markets were originating from the vicinity (< c. 100 km) of the surveyed sites, and thus could be used as proxies of “populations” within the large-scale geographic framework (c. 1,000 km North-South) of our study. Free consent from hunters and market sellers was obtained before collecting samples. We relied on an opportunistic sampling strategy (see Olayemi et al., 2011), without financial incentives. Sample types consisted of fresh skin, tissue (muscle) and tongue taken from dead animals. In each case, samples were taken beneath the tissue surface, so to prevent any potential exogenous contamination.

Genomic DNA extraction was performed using the NucleoSpin® Tissue Kit (Macherey- Nagel, Hoerdt, France), following manufacturer’s recommendations. Final elution step was repeated twice in 50 µl elution buffer to increase DNA yield. DNA concentration was quantified using the NanoDrop 1000 Spectrophotometer (ThermoFisher Scientific, Illkirch-Graffenstaden, France).

We amplified a mitochondrial fragment of 402 bp from cytochrome *b* (cyt *b*) able to differentiate among the six WBP lineages, following Gaubert et al. (2016). PCR products were sequenced at Genoscreen (https://www.genoscreen.fr/en; Lille, France) and Macrogen Europe (https://dna.macrogen-europe.com/en; Amsterdam, the Netherlands). Sequences were manually aligned in BioEdit v7.0.5 (Hall, 1999). All the sequences produced in this study were deposited in Genbank under accession numbers OP524750 - OP525226.

We amplified 20 microsatellite markers developed from the genome of WBP using four multiplexes (4 to 6 loci) following Aguillon et al. (2020). PCR products were run on an ABI 3730 DNA Analyser (Thermo Fisher Scientific) at Genoscreen and GeT-PlaGe (https://get.genotoul.fr/; INRAE, Toulouse, France).

### Data analysis – Cytochrome *b*

#### Clustering analysis

We used cyt *b* to phylogenetically assign the collected samples within the framework of the six mitochondrial lineages as previously defined (Gaubert et al., 2016). For that purpose, we combined our samples from Cameroon (N = 525), Equatorial Guinea (N = 15), Bioko Island (N = 1), Gabon (N = 18) and European seizures (N = 20) to the cyt *b* sequences available for WBP at the time of the study (N = 47). A clustering distance tree was obtained with MEGA-X v10.2.2 (Kumar et al., 2018) using Neighbor Joining and the Kimura 2-parameter model (Kimura, 1980). Node support was assessed through 500 bootstrap replicates (Felsenstein, 1985).

#### Genetic diversity and structure

Based on the new cyt *b* sequences that we produced, we reassessed the levels of genetic variation in WCA and Gab, after removing sequences with missing data (remaining N = 565). We used DnaSP v6.12 (Rozas et al., 2017) to compute haplotype number (*h*), haplotype diversity (*Hd*) and nucleotide diversity (*π*) (Supplementary Table S2 online). We mapped the geographic distribution of WCA and Gab haplotypes using ArcGIS 10.1 (Esri France). We ran Network v10.2.0.0 (Polzin & Daneshmand, 2008) to build a median-joining haplotype network with ε = 0, so to minimize the presence of alternative median networks.

#### Demographic history

Because of the low number of samples attributed to Gab (N = 13; see Results) and after removing sequences with missing data, we explored the demographic history of WCA only (N = 552). We performed mismatch analysis using Arlequin 3.5.2 (Excoffier & Lischer, 2010) to test for signatures of demographic and spatial expansions, by calculating the sum of squared deviations (SSD) between observed and expected distributions using 1,000 bootstrap replicates (Schneider & Excoffier, 1999).

Past variation in effective population size was assessed through a Bayesian skyline plot analysis in BEAST 2.6.3 (Bouckaert et al., 2019). We used the Bayesian Model Test option (bModelTest; Bouckaert et al., 2019) for site modeling, a coalescent Bayesian skyline prior and a relaxed clock (log normal-distributed). We fixed the mean mutation rate (*ucld* mean) to 0.026 substitution/site/Myrs with a normal distribution ranging from 0.010 to 0.045, as extracted from the posterior distribution values of the calibrated tree of Gaubert et al. (2016; data not shown). We ran 500.10^6^ MCMC iterations, with trees and model parameters sampled every 10,000 generations and a 10% burnin. Parameter convergence and Bayesian skyline plot reconstruction were, respectively, visualized and performed in Tracer 1.7.1 (Rambaut et al., 2018). Effective population size (N_e_) was estimated considering a generation time of 2 years in WBP (Zanvo et al., 2022).

### Data analysis – Microsatellite markers

#### Genetic diversity

We used Geneious 9.0.5 (Kearse et al., 2012) and the Microsatellites plugin (https://www.geneious.com/features/microsatellite-genotyping/) for allele scoring and genotype extraction. Only WCA and Gabon individuals with at least 75% of genotyping success were considered for the analyses (N = 558). The 75% threshold coincides with ≥ 15 microsatellite markers, which triples the minimum number of loci needed to discriminate among WCA individuals (Aguillon et al., 2020).

Detection of null alleles in the 20 loci was performed with Microcheker 2.2.3 (Van Oosterhout et al., 2004), assuming population at equilibrium, on the Foumbot population (N = 31). Deviation from Hardy-Weinberg Equilibrum (HWE) was calculated in GenAlEx 6.5 (Peakall & Smouse, 2012). Linkage Deseliquilibrum (LD) was assessed using FSTAT 2.9.4 (Goudet, 2003) after 1,000 randomizations (*P-value* ≥ 0.0000526316). The Bonferroni correction was applied to null hypothesis testing in those three analyses.

Genetic diversity was assessed through the number of alleles (Na), observed (*Ho*), expected (*He*) and unbiased expected (*uHe*) heterozygosity (in GenAlEx), allelic richness (A_R_) and deficit of heterozygotes (*F_IS_*) (in FSTAT), as mean values per population (see below).

#### Genetic structure

Global genetic variance among all genotyped individuals was assessed through pairwise population matrix unbiased genetic distances, and was visualized in GenAlEx using Principal Coordinates Analysis (PCoA).

For downstream population-based analyses, we defined the following 11 populations based on geographic proximity and biogeographic continuum among forests: Bayomen (Bayomen and Bangong [Bafia forest]; N = 5), Mt Cam (Manyemen and Bayib-Assibong [forest around National Park of Mt Cameroon]; N = 6), Eseka (Eseka, Mamb, Lolodorf, Bipindi and Eseka [Bassa’s forest] N = 22), GES (forests of southern Equatorial Guinea; N = 12), PNCM (Campo, Ma’an [forest of National Park of Campo Ma’an] and northern Equatorial Guinea; N = 11), Foumbot (Foumbot [Noun/Tikar forests]; N = 36), Yabassi ([Ebo Forest]; N = 9), Manengole ([Manengumba Bakossi Forest]; N = 19), Abong Mbang (Lomié [forests of East Cameroon]; N = 15), Sangmelima (Sangmelima and Djoum [forests around Dja Biosphere Reserve]; N = 45), and Gab (North and South of Gabon; N = 5). Samples from Yokadouma (N = 3), Ekombitie (N = 1), Esse (N = 2), Akonolinga (N = 1), Nditam (N = 3), Yokadouma (N = 2) and Bioko Isl. (N = 1) (see Supplementary Table S1 online) were excluded because not proximate enough to any site to be merged and reach a sufficient number of samples.

We used STRUCTURE 2.3.4 (Pritchard et al., 2009) to conduct a clustering analysis, following a two-step dataset refinement procedure, where (i) all the individuals from the study area were included (WCA + Gab; N = 558), and (ii) only the WCA individuals were included (WCA; N = 181), discarding large urban markets and cyto-nuclear hybrids or admixed individuals (who with at least 20% of shared ancestry). The first scheme allowed screening the global genetic structure of WBP at the largest study scale. The second scheme aimed at assessing the genetic structure among WCA reference populations. The first and second schemes were run with the LOCPRIOR model, notably to manage the sampling bias among lineages (first scheme) and populations (second scheme; Hubisz et al., 2009)). We performed for each scheme 20 independent runs for K = 2 to 10 with 10^5^ Markov chain Monte Carlo (MCMC) iterations (burnin = 10^4^), assuming admixture and uncorrelated (first scheme) or correlated (second scheme) allele frequencies. We used STRUCTURE HARVESTER 0.6.94 (Earl & vonHoldt, 2012) to estimate the most likely number of populations (K) using the ɅK method (Evanno et al., 2005). Graphic outputs from independent runs were summarized with CLUMPP (Jakobsson & Rosenberg, 2007), and CLUMPAK server (http://clumpak.tau.ac.il/) was used to generate the bar plots of population assignment.

We also inferred the most likely number of geographic populations using WCA georeferenced individuals, through the *Geneland* package (Guillot et al., 2005) in R 4.0.5 (R Core Team, 2021). We followed Coulon et al. (2006) by first allowing K to vary from 1 to 20 and launched five runs of 5.10^5^ MCMC iterations (thinning = 500; burnin = 500) under the frequency-correlated model and 1 km of uncertainty for spatial coordinates. Second, we fixed the number of estimated populations to K = 10-13 after the results of the first analysis [highest mean posterior densities for: K = 10 (0.21), K = 11 (0.30), K = 12 (0.26) and K = 13 (0.15)], and performed 20 independent runs with the same parameters. In the final analysis, the stability of individual assignment to populations was assessed among the best five runs (*i.e.* with the highest posterior probabilities). We used 500 × 500 pixels to map the posterior probabilities of population membership.

Pairwise differentiation (*F_ST_*) among the 11 populations as defined above was computed in Arlequin 3.5 (Excoffier & Lischer, 2010). Significant differentiation values (P ≤ 0.05) were estimated using 10^5^ MCMC iteration chains and 10,000 dememorization steps.

We tested isolation-by-distance (IBD) within WCA individuals and among WCA populations through the R package *pegas* (Paradis et al., 2004) using a Mantel test. The significance of the correlation (*r*) between individual-based genetic (Edward’s) and geographic (Euclidean) distances was estimated through 10,000 permutations.

#### Demographic history

We tested for the signature of demographic bottlenecks in WCA (excluding hybrid and admixed individuals) using both the Single Mutation Model (SMM) and the Two Phase Model (TPM) in BOTTLENECK 1.2.02 (Piry et al., 1999). We applied the Wilcoxon sign-rank test to estimate heterozygote excess/deficit using 10,000 replications.

We assessed temporal changes in the effective population size of WCA through an approximate-likelihood approach implemented in the R package *VarEff* (Nikolic & Chevalet, 2014). We followed Zanvo et al. (2022) by fixing a generation time of 2 yrs and a microsatellites mutation rate of 5.10^-4^ per generations (Schlotterer, 2000). We ran the analysis with three mutation models (single, geometric and two phases), using 10,000 MCMC batches (length = 1, thinned every 100 batches; JMAX = 3). The first 10,000 batches were discarded as burnin.

#### Tracing the pangolin trade

The discriminative power of our 20 microsatellite markers among WBP individuals was evaluated after three different approaches: (i) counting the number of identical genotypes among samples using the Multilocus tagging option in GenAlEx (suboption Matches), (ii) computing the probability of encountering the same genotype more than once by chance using the R package *poppr* (method=single; Kamvar et al. (2014), and (iii) calculating the indices of unbiased probability of identity and probability of identity among siblings (uPI and PIsibs) using Gimlet 1.3.3 (Valière, 2002).

Because we could not find a deep level of genetic structure within WCA, we followed the protocol based on private allele frequency rarefaction described by Zanvo et al. (2022) to trace the trade of WBP in our study area. We used the generalized rarefaction approach implemented in ADZE (Szpiech et al., 2008) to compute private allele frequencies (paf) among various combinations of populations from the 10 source populations as defined above (removing Gab). Because the original scheme of 10 populations yielded the greatest inferred values of paf (data not shown), we graphed the paf rarefaction curves (sample size rarefied from 2 to 5, 5 being the lowest number of individuals among populations) for each locus in these 10 populations. Only loci for which paf were > 40% and reached a plateau or showed an increasing trend at N = 5 for a given population were considered as potentially useful for tracing the origin of WBP found in the urban markets. We then cross-filtered these results with the private alleles observed in the actual populations (using GenAlEx), and eventually retained the loci that showed both high potential for tracing (ADZE output) and observed private alleles (GenAlEx output) for the given populations. As a final step, we manually screened the private alleles present in WBP sampled from the Cameroonian large urban markets (Douala and Yaoundé) and international seizures to trace their geographical origins.

## Results

### Mitochondrial DNA

The Neighbor joining (NJ) phylogenetic tree based on 626 cytochrome *b* (cyt *b*) sequences recovered the six WBP geographic lineages, including Western Africa, Ghana, Dahomey Gap, WCA, Gab and Central Africa (Supplementary Fig. S2 online). A total of 569 individuals from Cameroon, Equatorial Guinea and northern Gabon (Bitam) clustered within WCA, whereas ten samples from southern and northern Gabon and three samples from central-south Cameroon (Yaoundé and Abong Mbang) clustered within Gab (Fig. 1).

**Figure 1.**
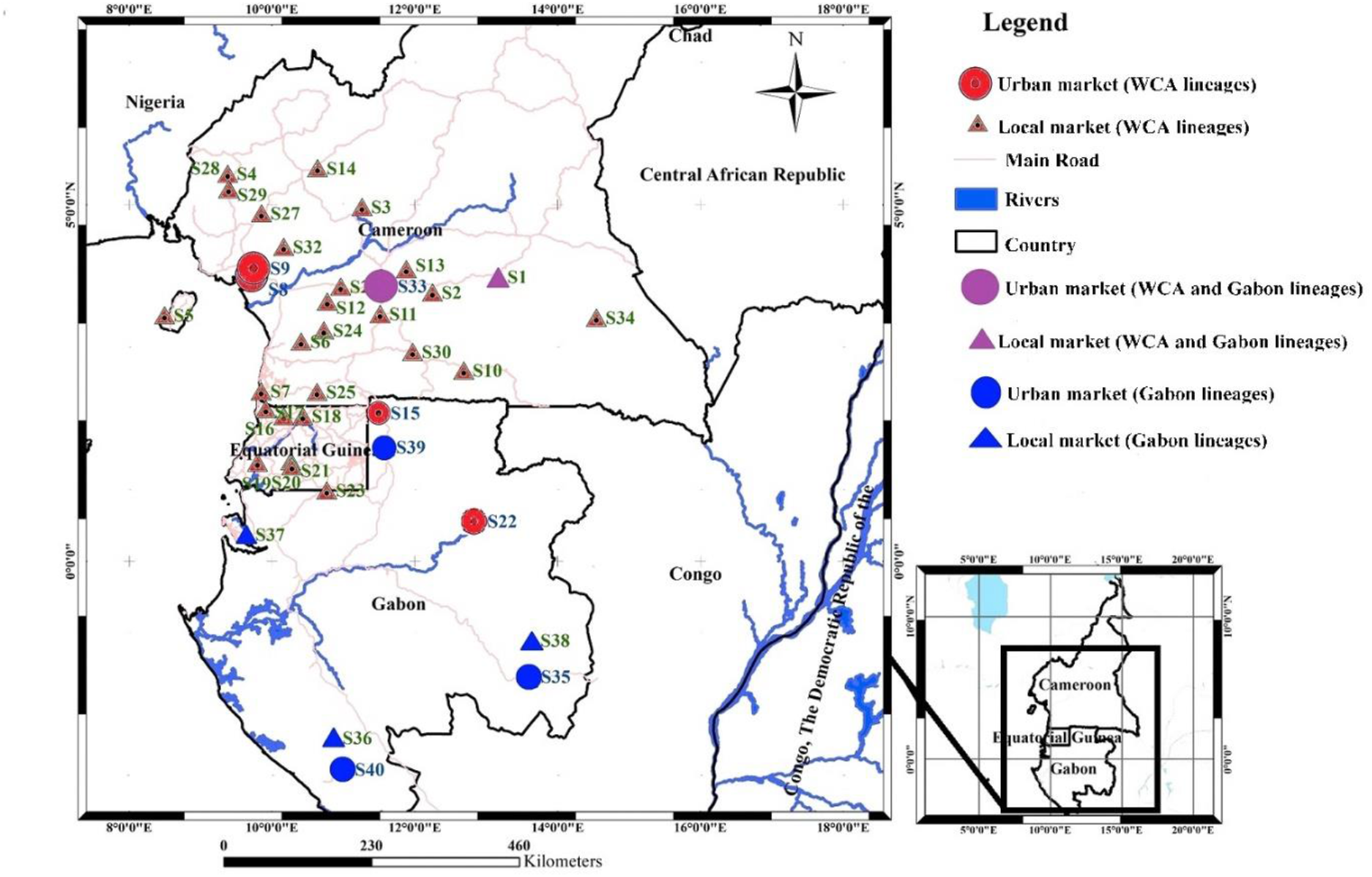
Reassessed distribution of the Western Central Africa and Gabon mitochondrial lineages in white-bellied pangolins. Lineage assignment was based on NJ tree clustering (see Results and Supplementary Fig. S2). Red = Western Central Africa lineage; Blue = Gabon lineage; Purple = sites were the two lineages co-occur.

S1-Abong Mbang; S2-Akonolinga; S3-Bangong; S4-Bayib Assibong; S5-Bioko; S6-Bipindi; S7-Campo; S8-Douala Central Market; S9-Douala Dakat Market; S10-Djoum; S11-Ekombitié; S12-Eseka; S13-Esse; S14-Foumbot; S15-Bitam; S16-Anguma; S17-Bongoro; S18-Ebenguan; S19-Emangos; S20-Misergue; S21-Taguete; S22-Makokou; S23-Medouneu; S24-Lolodorf; S25-Maan; S26-Mamb; S27-Manengole; S28-Manyemen; S29-Nditam; S30-Sangmelima; S31-Seizure; S32-Yabassi; S33-Yaoundé Nkolndongo Market; S34-Yokadouma; S35- Franceville; S36-Mocabe; S37-Nkoltang; S38-Okoumbi; S39-Oyem; S40-Tchibanga.

A total of 67 and seven haplotypes were identified in WCA and Gab lineages, respectively. WCA and Gab were differentiated by 23 mutations in the median-joining haplotype network, whereas no particular geographic structure could be observed within the two lineages (Supplementary Fig. S3 online). Three haplotypes were dominant in WCA, with H2 (28%) and H3 (25%) located only in Cameroon, and H5 (17%) in Cameroon and Bioko Island (Supplementary Fig. S4 online). In Gab, the dominant haplotype (H70; 50%) was distributed from northern to southeastern Gabon. The two Douala markets recruited 8-17 haplotypes (dominated by H5 and H2 at 27% and 23%, respectively), while the Yaoundé market was characterized by 35 haplotypes (dominated by H3 at 32%). Haplotype (*Hd*) and nucleotide (π) diversities were respectively 0.830 and 0.010 in WCA, and 0.897 and 0.019 in Gab (Supplementary Table S2 online).

Based on mismatch analysis, the hypotheses of spatial and sudden demographic expansions could not be rejected (P(Sim. Rag. ≥ Obs. Rag.) = 0.559 and P(Sim. Rag. ≥ Obs. Rag.) = 0.424 for WCA (see Supplementary Fig. S5 online).

Bayesian skyline plots showed a c. 22-fold increase in the median effective population size of WCA from N_e0_ = c. 1900 (HPD 95% = c. 750 – 4,200) to N_e1_ = c. 41,500 (HPD 95% = c. 11,000 – 206,000), abruptly starting c. 275 kya (Fig. 2).

**Figure 2.**
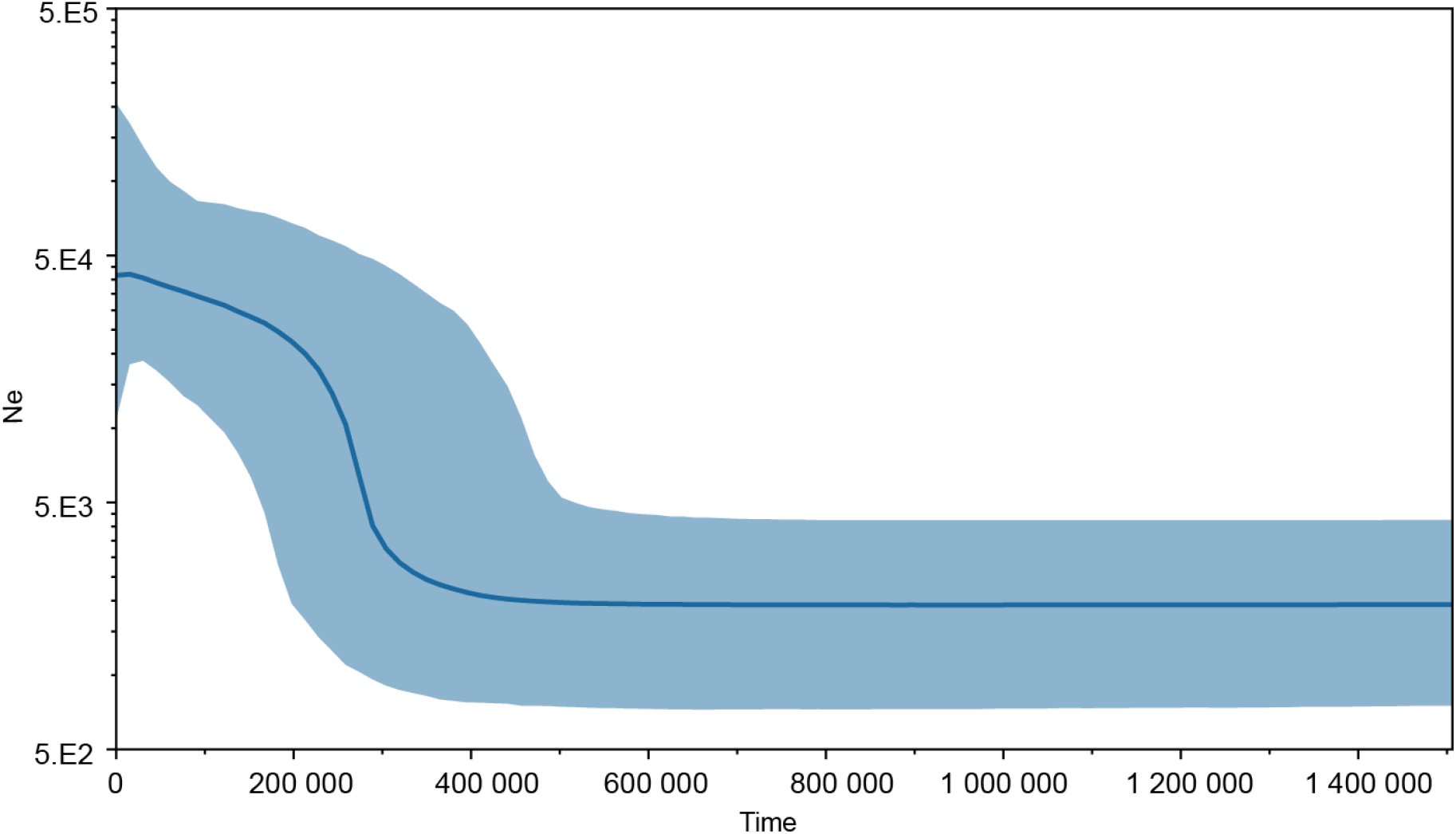
Bayesian skyline plots showing the demographic history of white-bellied pangolins from the Western Central Africa lineage. Y-axis = Effective population size (N_e_); X-axis = Time in years before present.

### Microsatellite markers

#### Genetic diversity

In the reference population (Foumbot) that we used to assess locus equilibrium parameters (Foumbot), we detected potential evidence for eight loci affected by null alleles (Supplementary Table S3 online) and significant deviation of genotypic frequencies from those expected under HWE at four loci (Supplementary Table S4 online). No linkage disequilibrium between pairs of loci could be identified.

Among populations, mean number of alleles per locus (Na) was 6.4 and ranged from 4.0 (Bayomen) to 12.7 (Yaoundé market). The overall mean observed heterozygosity (*Ho*) was

0.579 (0.534 - 0.610) and average expected heterozygosity (*He*) was 0.664 (0.593 - 0.710) (Table 1). Overall, inbreeding coefficient (F_IS_) values were positive (F_IS_ = 0.168) and ranged from 0.112 in GES to 0.217 in Eseka. Seven F_IS_ values were significant. The mean allelic richness (A_R_) was 4.093 and was the highest in airport seizure (5.285) and urban markets (Table 1). Numbers of private alleles were the greatest for Gabon (15) and Sangmelima (11) when the urban bushmeat markets from Yaoundé and Douala were not included. When included, Yaoundé showed the highest number of private alleles (42). Observed frequencies of private alleles were the greatest for Gabon (0.922-0.932) whether urban bushmeat markets were included or not. Frequencies of private alleles were low for the Douala (0.111) and Yaoundé (0.141) markets.

**Table 1.** Genetic diversity across reference populations, urban bushmeat markets and seizures of white-bellied pangolins from Cameroon, Equatorial Guinea and Gabon, as estimated from 20 microsatellites loci.

| Populations | N | Na | Ho | He | uHe | F<sub>IS</sub> | A<sub>R</sub> | No. private alleles (without urban markets) | No. private alleles (with urban markets) | Private allele frequency (without urban markets) | Private allele frequency (with urban markets) |
| --- | --- | --- | --- | --- | --- | --- | --- | --- | --- | --- | --- |
| Abong Mbang | 13 | 6.200 | 0.571 | 0.685 | 0.713 | 0.207 | 3.997 | 5 | 1 | 0.320 | 0.196 |
| Bayomen | 5 | 4.000 | 0.590 | 0.593 | 0.659 | **0.116** | 3.645 | 0 | 0 | 0.200 | 0.140 |
| Eseka | 23 | 7.150 | 0.548 | 0.687 | 0.703 | **0.217** | 3.890 | 4 | 2 | 0.217 | 0.134 |
| Foumbot | 36 | 7.400 | 0.561 | 0.657 | 0.666 | **0.159** | 3.664 | 4 | 1 | 0.162 | 0.097 |
| GES | 12 | 5.850 | 0.610 | 0.657 | 0.685 | 0.112 | 3.812 | 5 | 2 | 0.211 | 0.147 |
| Manengole | 19 | 6.250 | 0.608 | 0.671 | 0.690 | **0.122** | 3.786 | 2 | 1 | 0.186 | 0.113 |
| Mt Cam | 6 | 4.500 | 0.609 | 0.628 | 0.690 | 0.130 | 3.850 | 2 | 1 | 0.240 | 0.139 |
| PNCM | 11 | 5.200 | 0.580 | 0.650 | 0.682 | **0.149** | 3.625 | 1 | 1 | 0.127 | 0.071 |
| Sangmelima | 45 | 8.350 | 0.587 | 0.704 | 0.712 | **0.170** | 3.955 | 11 | 4 | 0.244 | 0.160 |
| Yabassi | 9 | 5.700 | 0.595 | 0.647 | 0.676 | 0.138 | 3.632 | 2 | 1 | 0.144 | 0.073 |
| Gabon | 5 | 4.850 | 0.575 | 0.638 | 0.710 | **0.210** | 4.332 | 15 | 8 | 0.932 | 0.922 |
| Douala markets | 84 | 9.300 | 0.546 | 0.687 | 0.691 | - | 4.857 | - | 8 | - | 0.111 |
| Yaoundé market | 248 | 12.700 | 0.534 | 0.710 | 0.712 | - | 4.979 | - | 42 | - | 0.141 |
| Seizure (Europe) | 10 | 6.000 | 0.591 | 0.680 | 0.716 | - | 5.284 | - | 1 | - | 0.166 |

Populations are based on geographic proximity and biogeographic continuum among forests, lineage or seizure/selling places (urban markets): Gabon (corresponding to Gab lineage), Bayomen (Bayomen and Bangong [Bafia forest]), Mt Cam (Manyemen and Bayib-Assibong [forest around National Park of Mt Cameroon]), Eseka (Eseka, Mamb, Lolodorf, Bipindi and Eseka [Bassa’s forest]), GES (forests of southern Equatorial Guinea), PNCM (Campo, Ma’an [forest of National Park of Campo Ma’an] and northern Equatorial Guinea), Foumbot (Foumbot [Noun/Tikar forests]), Yabassi (Ebo Forest), Manengole (Manengumba Bakossi Forest), Abong Mbang (Lomié [forests of East Cameroon]), and Sangmelima (Sangmelima and Djoum [forests around Dja Biosphere Reserve]).

N: number of samples; Na: number of alleles; Ho: observed heterozygosity; He: expected heterozygosity; uHe: unbiased expected heterozygosity; F_IS_: inbreeding coefficient (values in bold are significant); A_R_: allelic richness.

#### Genetic structure

The PCoA did not show any clear structuring of the genetic space within WBP from WCA and Gab lineages (Supplementary Figure S6 online).

Clustering analysis with STRUCTURE, within all individuals and with prior information on lineage detected K = 2 as the most likely number of clusters (Fig. 3A). The two clusters did not totally correspond to the mitochondrial geographic assignment of the individuals regarding WCA and Gab lineages (Fig. 3A). In this case, 98% of all WBP individuals of WCA clustered into one population. Five out of eight mtDNA-assigned to Gab individuals clustered into a single group (Gab lineage). Three individuals (Y110, Ab10 and Ab2) from Yaoundé and Abong Mbang in Cameroon that were mtDNA-assigned to Gab clustered with WCA. Three individuals from Yaoundé and Douala (Y360, Y101 and DlaB103) that were mtDNA-assigned to WCA clustered with Gab. One individual from Yokadouma (Ou4) in Cameroon showed an admixed pattern between WCA and Gab (c. 60/40% assignment, respectively). From K = 3, no specific population structure was found (Supplementary Fig. S7 online).

**Figure 3.**
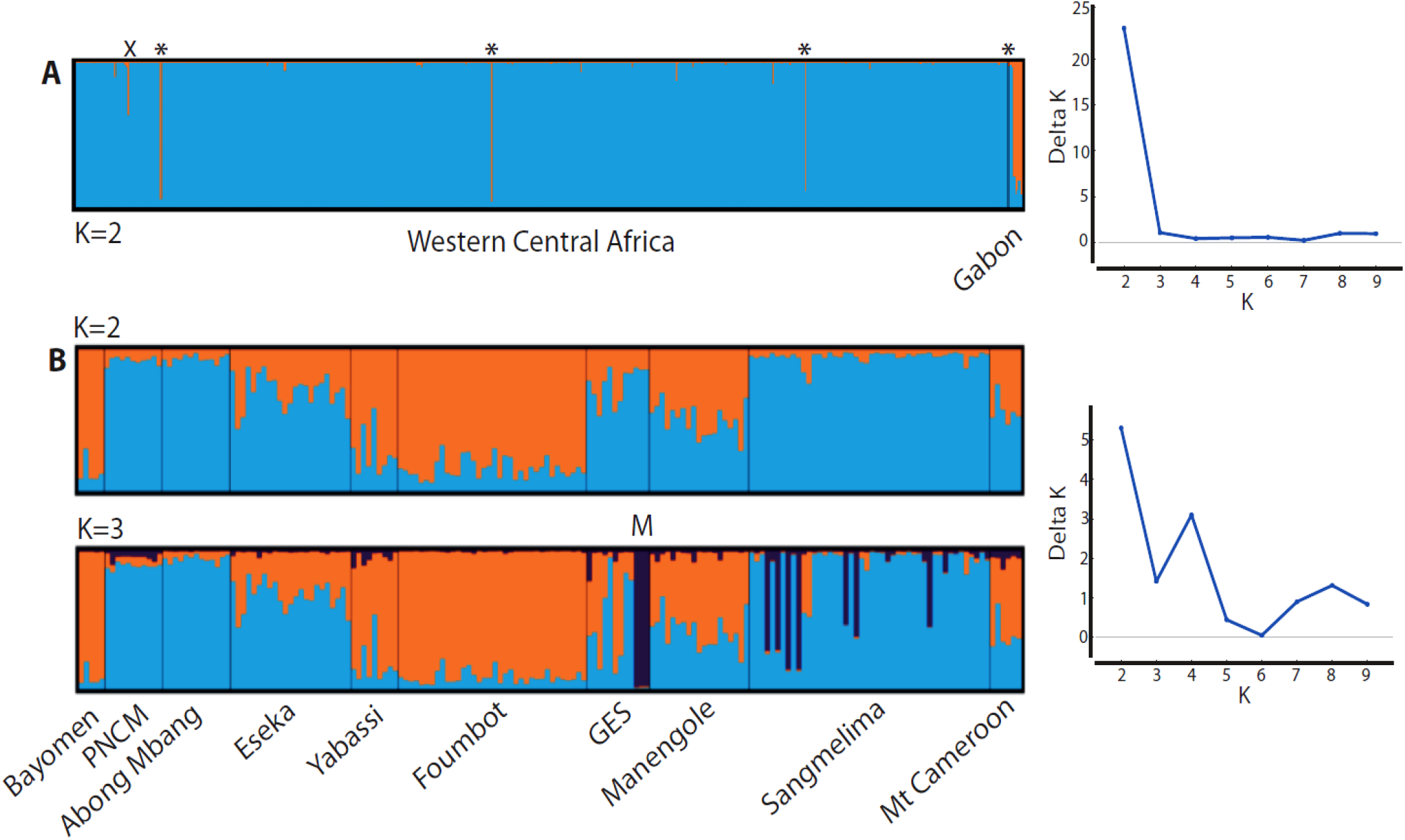
Assignment plots among individuals of white-bellied pangolins from western central Africa as obtained with STRUCTURE (left) and most likely number of population clusters (K = 2) following the ɅK method (right).

Within WCA and among reference populations, K = 2 was the most likely number of clusters (Fig. 3B). Two ‘pure’ genetic clusters regrouped Bayomen, Yabassi, and Foumbot from western and central Cameroon on one side, and PNCM, Abong Mbang and Sangmelima from North of Equatorial Guinea and southern / central Cameroon on the other side. Several reference populations from western, central and southern Cameroon, North of Gabon and South of Equatorial Guinea were admixed (Eseka, GES, Manengole and Mt Cam), and did not correspond to coherent geographic delineations (Fig. 3B). At K = 3, the four individuals from Medouneu (northern Gabon; GES population) clustered into an exclusive group. Seven individuals from Sangmelima showed full to admixed assignment to the Medouneu cluster. From K = 4, no further coherent geographic structure was found (Supplementary Fig. S8 online).

Each individual is represented by a vertical bar. A- including all the samples from urban markets, seizures and reference populations (N = 558), sorted according to their mtDNA-based lineage assignment (WCA *vs*. Gab); asterisks indicate cyto-nuclear hybrids; X is an admixed individual (> 20% of genomic ancestry shared with Gab). B- only retaining the samples from 10 reference populations (N = 181); M = cluster from Medouneu (northern Gabon).

Spatial genetic clustering analysis (Geneland) within WCA did not allow to geographically delineate consistent populations. Optimal number of populations varied between 10 and 13 clusters (Supplementary Fig. S9 online), with 11 as the most probable number of clusters according to the second-step iterative run (Supplementary Fig. S9 online). Assignment of individuals to populations among the best five runs was greatly unstable (data not shown).

F_ST_ values were significantly different (*p ≤ 0.001*) between all WCA populations and Gab (from 0.184 to 0.225; Table S5). Pairwise differentiation between WCA populations was low, ranging from 0.006 (PNCM - Sangmelima) to 0.048 (Bayomen - Southern Equatorial Guinea).

We detected a significant IBD pattern among WBP individuals across the studied range (*r*= 0.099, *p* = 0.001), whereas there was no significant effect among populations (*r* = 0.298, *p* = 0.094; Supplementary Fig. S10 online).

#### Demographic history

We detected a significant signal of bottleneck in WCA with the S.M.M. model (Wilcoxon sign-rank test; *p* < 0.05), whereas the T.P.M. model did not (*p* = 0.061).

The assessment of the historical demography of WCA with *VarEff* showed that effective population size (*Ne*) drastically declined –regardless of the models– over the last c. 9,840-6,140 years (4920-3070 generations; Fig. 4). Harmonic means estimated a 74.1-82.5% *Ne* reduction, from 7,773-13,826 (ancestral *Ne*) to 2,311-5,027 (contemporaneous *Ne*).

**Figure 4.**
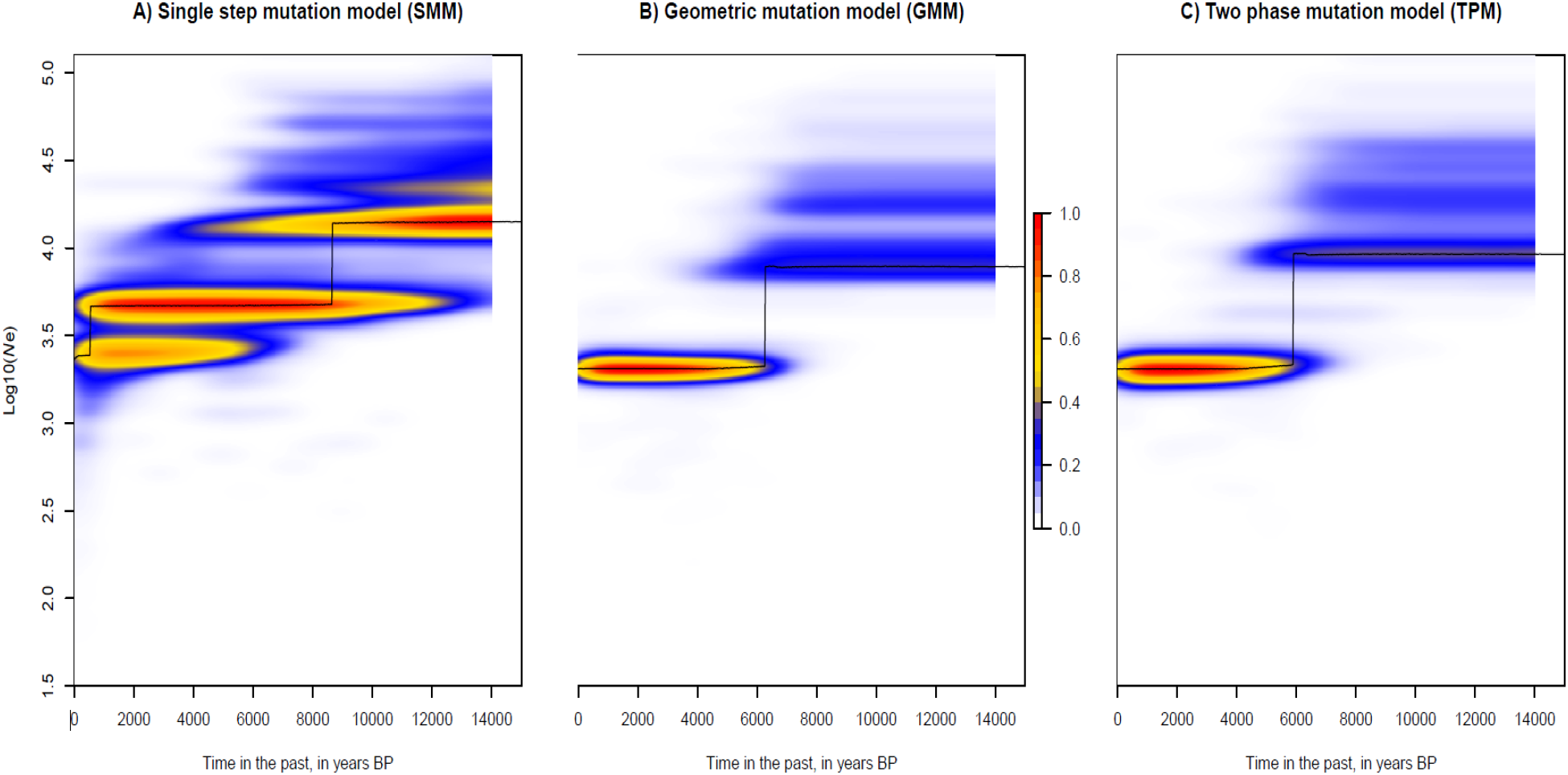
Temporal change in effective population size (*Ne*) of white-bellied pangolins from Western Central Africa, as estimated using *VarEff* under three different mutation models. Harmonic means (black line) and kernel density (color scale) of *Ne* posterior distributions are given in years BP.

#### Tracing the pangolin trade

A total of 535 (96%) of 557 WBP samples had unique genotypes. Eleven pairs of samples from Djoum, Manengole, Yaoundé and Medouneu shared a same genotype. The null hypothesis of encountering the same genotypes more than once by chance was rejected for all pairs of individuals (*p* < 0.0001). Unbiased probability of identity (uPI) and probability of identity among siblings (PIsibs) were both very low (uPI = 0.00125±0.00538; PIsibs = 0.0236±0.07236; Supplementary Fig. S11 online).

Seven of a total of 11 loci that presented appropriate private allele signatures for one or several populations using private allele rarefaction, could be related to 18 private alleles observed in the reference populations (WCA; Supplementary Table S6 online). On this basis, we could trace a total of 47 (12.8%) individuals sold as part of the bushmeat trade in the study region (Supplementary Table S7 online) to seven reference populations (see Fig. 5). Thirty-five WBP from the Yaoundé market were harvested from Sangmelima, Eseka, Mt Cameroon, Abong Mbang, Manengole and South of Equatorial Guinea. Ten individuals sold in Douala markets came from Abong Mbang, Sangmelima and Yabassi. Two pangolins seized at Brussels airport were traced to South of Equatorial Guinea and Abong Mbang.

**Figure 5.**
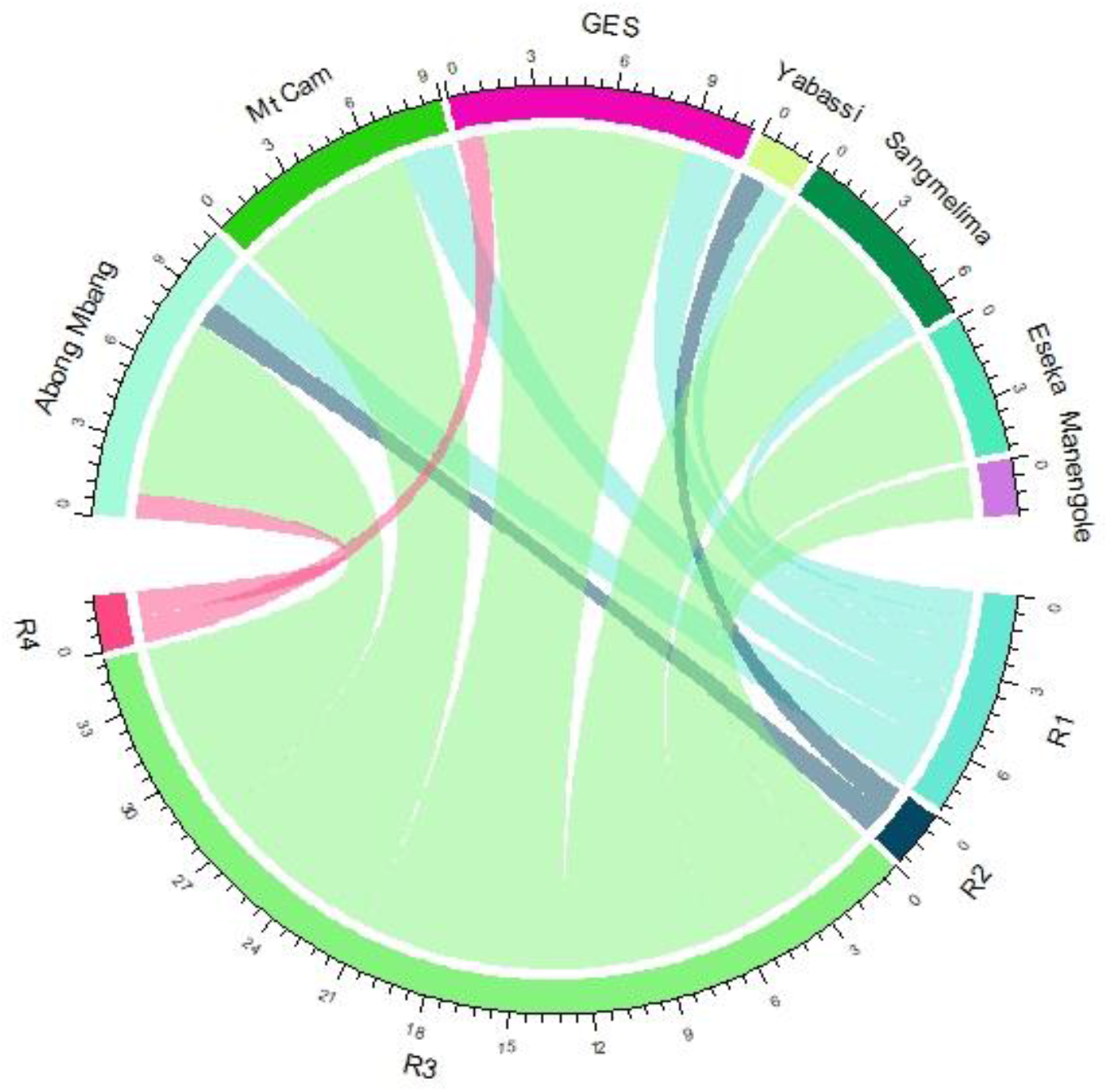
Chord diagram tracing the geographic origin of white-bellied pangolins from urban bushmeat markets and seizures. R1 = Douala Dacat market; R2 = Douala Central market; R3 = Yaoundé Nkolndongo market; R4 = Seizures at Brussels airport.

## Discussion

### White-bellied pangolins from western central Africa: range-overlapping lineages and weak geographic structure

Our exhaustive sample set across three bordering countries, representing 565 newly collected samples of WBP from Cameroon, Equatorial Guinea and Gabon, allowed refining the lineage distribution of WBP from western central Africa. We show that the two mitochondrial lineages described from the study region are overlapping in ranges, with Gab being present all over Gabon and reaching southern Cameroon, while the southern extension of WCA reaches northern Gabon. Thus, our study clarifies the geographic distribution of the two lineages in western central Africa as previously based on a reduced number of samples (Gaubert et al., 2016), and suggests that Gab is not endemic to Gabon. The three individuals representing Gab lineage in Cameroon were found in the Yaoundé market and in the reference population of Abong Mbang. The latter corresponds to a local market that is off the main road network connecting northern Gabon to southern Cameroon (Randolph et al., 2022). The involved market stakeholder indicated that the two pangolins originated from the Lomié forest (c. 100 km from the selling site), thus suggesting that the Gab lineage naturally occurs in Cameroon.

Microsatellites genotyping showed that WCA and Gab corresponded to differentiated evolutionary units (mean F_ST_=0.192), in line with previously published nDNA sequencing data (Gaubert et al., 2016) and with the mtDNA pattern found in the present study. However, because the three mtDNA-assigned Gab individuals from Cameroon were cyto-nuclear hybrids (i.e., assigned to the WCA cluster using microsatellites), ancient gene flow between the two lineages might have occurred and is blurring their geographic delineation, notably in their supposed range of co-occurrence. All the more since an admixed individual from Yokadouma, in south-eastern Cameroon, provides evidence for ongoing (but limited) introgression between the two lineages. Further sampling in forests from southern Cameroon and northern Gabon will have to be conducted in order to achieve a more comprehensive picture of the respective ranges of the WCA and Gab genetic entities, and their potential zones of admixture.

Despite the use of hypervariable microsatellite markers and a fair level of genetic diversity observed across populations (Aguillon et al., 2020; this study) along the c. 1,000 km extension of our study zone, genetic structure and differentiation within WCA was weak. Clustering analysis (Structure) was the sole capable of detecting population structure, using the available signal of geographic structure based on prior delineation of populations (r < 1; data not shown). However, the two ‘pure’ genetic clusters did not correspond to anything coherent in the geographical space, with a first cluster located strictly south of the Sanaga river (PNCM + Abong Mbang + Sangmelima) and the other spread on both sides of the river (Bayomen [north] *vs.* Yabassi and Foumbot [south]), while admixed individuals were present north and south of the river (Manengole and Mt Cam [north] *vs.* Eseka and GES [south]).

At K=3, WBP from northern Gabon (Medouneu), which corresponds to the southernmost limit of WCA, were identified as a distinct population cluster. Because Medouneu is located c. 50 km from Monts de Cristal, a putative Pleistocene rainforest refugium (Maley, 1996), pangolins from this area may hold the signature of a relict population, as previously observed in mammals and plants (e.g., Anthony et al., 2007; Faye et al., 2016). However, the lack of population signature in other areas of putative refugia such as southern Gabon and western Cameroon (Maley, 1996; both sampled in our study) questions this hypothesis; all the more since Medouneu is located at the bottom end of the North-South IBD pattern that we observed in WCA (see below), which could be the cause of such apparent clustering (see Perez et al., 2018). The refugium hypothesis is also challenged by the presence of Medouneu-related individuals (pure or admixed) further north in southern Cameroon (Sangmelima), suggesting a more extended range for this population.

The lack of genetic structure and population differentiation across WCA is to be interpreted in the light of the IBD pattern that we detected among WBP individuals. Although our study did not quantify dispersal rate and range, the pattern that we observed fits a stepwise migration model where dispersal distances are large enough to lead to near-panmictic genetic patterns (Kinitz et al., 2013). Our results confirm the non-influential role of the Sanaga river on the biogeography and population delimitation of several African taxa (see Fuchs & Bowie, 2015), although its role as a limiting –i.e., not prohibitive– factor to gene flow (e.g., Gonder et al., 2011) was not tested here. They also suggest that there is no specific barrier to gene flow for WBP in the study zone (WCA), in line with the lack of structure found in another lineage of WBP investigated with the same markers (Dahomey Gap; Zanvo et al., 2022). Our population genetic study again provides evidence for unsuspected dispersal capacities in WBP, which is also coherent with a recent, spatial and/or demographic expansion across the WCA range following forest post-glacial expansion during the Late Pleistocene (e.g., Piñeiro et al., 2017), as suggested by the mtDNA data. Despite habitat fragmentation and hunting activities occurring in the study zone (Din Dipita et al., 2022; Mahmoud et al., 2020; Ordway et al., 2019), dispersal may be maintained given the capacity of the species to cross and occur in anthropized habitats (Akpona et al., 2008; Jansen et al., 2020) and the still prevalent, large forested areas available (Gillet et al., 2016). Field surveys are direly needed to document the dispersal behavior of WBP, notably during their early phase of supposed vagrancy (Gaubert, 2011).

The only individual from Bioko Island that we could sequence and genotype did not show any genetic differentiation from the continent. Thus, the question of whether WBP sold on Bioko Isl. are imported from the continent (Cameroon) or not cannot be answered here (Ingram et al., 2019), and as a corollary, that of a surviving population endemic to the island. Additional genetic surveys in Bioko Isl., both from bushmeat markets and field sites, will have to be conducted in order to assess the status of WBP on the island, an urgent matter of conservation as island populations are often subject to rapid genetic drift, accumulation of deleterious variants and inbreeding (Fry et al., 2020; Keller & Waller, 2002; Robinson et al., 2016).

### High level of genetic diversity and Wahlund effect in the context of a recent demographic decline

Overall, populations from the WCA and Gab lineages were characterized by the highest levels of genetic diversity described in WBP. Such information is important, as high genetic diversity is expected to promote population fitness and ensure adaptability to environmental change (Habel & Schmitt, 2012). MtDNA nucleotide diversity was 1.7-9.5 times greater for WCA (largest sampling effort; N > 550) and Gab (reduced sampling effort, similar to other lineages; N = 13) compared to the other lineages. Mean allelic richness in WCA was also higher than in the Dahomey Gap lineage (*A_R_* = 10.79 *vs*. 4.27; minimum sample size = 169; Zanvo et al., 2022), the only comparable study available.

The generally significant deficit of heterozygotes observed among populations (F_IS_ = 0.112-0.217) could lead to the conclusion that WBP populations from WCA are inbred. However, inbreeding would come in contradiction with the fairly high levels of heterozygosity observed among populations (mean *Ho* = 0.579; mean *He* = 0.664) and the weak geographic structure observed across WCA (see Wright, 1969). Because population differentiation (F_ST_) across WCA was low whereas F_IS_ values were high, our results fit the expectation of a Wahlund effect, i.e. where a same population regroups individuals from genetically differentiated entities (De Meeûs, 2018). Such signature could be reached through regular, long-range inter- connectivity among WBP populations, implying again unsuspected dispersal capacities of the species. However, given the knowledge gaps affecting the natural history of WBP and African pangolins in general (Heighton & Gaubert, 2021), we cannot discard the fact that some unknown breeding strategies of the species could participate to this Wahlund effect.

We identified an important decline in the effective population size of WCA (as mirrored by the bottleneck signature detected in one of our analyses) occurring c. 4,300-7,440 ya, a time of climatic stability favourable to rainforest cover and that predated a major period of savannah expansion and forest fragmentation in western central Africa (starting c. 4000 ya; Lézine et al., 2013; Ngomanda et al., 2009). Rather than a paleoclimatic influence, we hypothesize that the arising of the Ceramic Late Stone Age culture in the region from c. 7000 BP onward – characterized by bifacial macrolithic and polished stone tools (Grollemund et al., 2015) – could have constituted the technological change (from “Stone to Metal Age”; Lavachery, 2001) that made hunting more efficient (see Klein & Gruz-Uribe, 1996; Orijemie, 2022) and responsible for the decline of WBP. Such decline, although drastic (loss of c. -74.10-82.5% of the effective population size), places the effective populations size of WBP from WCA (2161±219) within the conservative threshold range of minimum viable population size (500–5000; Clabby, 2010; Reed et al., 2003). Because viable population size levels are paramount to maintaining acceptable levels of genetic diversity (Charlesworth, 2009), it will be important to monitor forest corridors among WBP populations and mitigate deforestation and illegal hunting activities in order to preserve the genetic diversity of pangolin populations in western central Africa.

### Yaoundé as a main domestic hub of an endemic, transnational trade of white-bellied pangolins in western central Africa

We showed that the 20 microsatellite loci specifically developed for WBP (Aguillon et al., 2020) provided the necessary power to confidently distinguish among all the WCA and Gab individuals. Only five microsatellite loci were needed to reach the conservative value of probability of identity < 0.01 (Waits et al., 2001), compared to the seven loci required for distinguishing pangolins from the Dahomey Gap (Zanvo et al., 2022). These results have important implications for the implementation of a convenient genetic tracing tool applied to the pangolin trade in western central Africa. Indeed, even a subset of our markers would be capable of estimating the exact number of individuals from scale seizures and trace the parallel trafficking networks of scale and body trade where same individuals can be dispatched (Ingram et al., 2019; Zanvo et al., 2021; Zhang et al., 2017). Microsatellites genotyping also allowed identifying eleven cases where a same pangolin had been sampled twice, illustrating the difficulties of relying on third parties such as trained local assistants who are part of the bushmeat chain (Din Dipita et al., 2022), but at the same time showing the usefulness of our markers to counter such sampling bias.

Although mtDNA was not resolutive enough to trace the trade at the local scale, we showed that the trade of WBP was endemic to western central Africa, involving both WCA and Gab lineages but excluding the other lineages described from central and West Africa (Gaubert et al., 2016). This is in line with the trade endemism also found for the Dahomey Gap lineage (Zanvo et al., 2022), and suggests that large, urban bushmeat markets (here, Douala and Yaoundé for the most sampled) are not necessarily hubs for long-range, international trade of WBP. Rather, they represent domestic hubs that source large volumes of WBP at the sub- regional level (i.e. country-wide and neighboring countries), but are not directly involved in global trade networks such as those identified from harbor or airport seizures in Africa and Asia (Emogor et al., 2021; Zhang et al., 2020).

The private allele frequency approach recently described (Zanvo et al., 2022) allowed us to fine-scale the geographic origins of 47 WBP sold from three urban bushmeat markets in Cameroon and seized at a European airport, representing c. 13% of the traceable individuals from WCA. Our results confirm that the Yaoundé market is the main national hub for the trade of bushmeat in Cameroon (Edderai & Dame, 2006; Randolph, 2016), recruiting among six of the seven geographic sources that were traced as part of this study. Sourcing occurs in the western, southern and eastern part of the country, but also in Equatorial Guinea, representing a distance network between c. 120 and 370 km. One geographic sourcing North of Douala and exclusive to the Yaoundé market, Manengolé, may provide evidence for transit of pangolins through Douala to Yaoundé, urban bushmeat markets in this case playing both a role of drop-off point and transit zone (Nguyen et al., 2021; Randolph et al., 2022). The two bushmeat markets from Douala showed contrasting spheres of influence, with Dacat being fed by a minimum of five geographic sources *contra* solely two sources for the Central market. Having said that, both showed a wide range of sourcing, respectively extending c. 600 and 500 km from Douala, beyond the maximal range delineated for Yaoundé. The two markets shared a unique supply source through the Yabassi population, situated c. 90 km north of Douala, in line with the recognized role of the unprotected Ebo forest as a feeding source of the Douala bushmeat market places (Din Dipita et al., 2022; Fuashi et al., 2019).

Eventually, we could trace the presence of WBP from the forests of southern Equatorial Guinea in Douala and Yaoundé markets, and Brussels airport (with flights originating from Cameroon). This suggests that transnational trade across neighbouring countries occurs in the study region, potentially following the main road network connecting Equatorial Guinea to Cameroon, and that Cameroon constitutes a major hub of this sub-regional trade through the large feeding network of its urban bushmeat markets (Ingram et al., 2019; https://cameroonvoice.com/news/2021/07/22/cameroun-trafiquants-de-pangolin-en-afrique-centrale-demasques/; this study). On the other hand, the likely occurrence of the Gab lineage and admixture between WCA and Gab or individuals from northern Gabon (Medouneu) in Cameroon blurred our tracing ability for a potential WBP trade occurring between Gabon and Cameroon, as previously mentioned in the literature (Ingram et al., 2018, 2019) and recent media covers (https://www.gabonmediatime.com/216-pointes-divoire-saisies-cameroun-provenant-gabon-reactions-de-lanpn-gouvernement-attendues/).

The number of haplotypes and number of private alleles were strikingly high in the Yaoundé market, supporting the hypothesis that Yaoundé is one of the main pangolin trade hubs of the study region. This was confirmed by the 12 different geographic sources mentioned by bushmeat vendors when asked about the origin of the pangolins sold in Yaoundé (N = 59; data not shown), including Sangmelima (and its surroundings), Mekin, Djoum, Akonolinga, Makenene, Bafia, Ma’an, Campo Ayos, Lomié, Yokadouma, Foumbot, Mbouda, Magba, Eseka, Mboumnyebel and Bankim. Such wide geographic network, obviously minimized by the genetic data set at hands, suggests a feeding network with a maximum sourcing distance of c. 600 km from the city, more in line with the market’s large network range expected from the literature (see above). The great number of private alleles detected for the Yaoundé market (42 *vs*. 0 to 8 in reference populations and Douala markets, and 15 in Gabon) could suggest that, despite our considerable sampling effort, major reference populations remain unsampled.

However, the relatively low private allele frequency calculated for Yaoundé supports the view that such high numbers of private alleles are an effect of sampling bias (> 250 samples). We believe that additional sampling of reference populations will have to be conducted across western central Africa to reach higher accuracy in the tracing of the WBP trade, although the theoretical numbers of sampled individuals for obtaining confident estimates of allele frequencies among populations for forensic purpose might be hardly achievable (see Chakraborty, 1992).

## Conclusion

Our study provides an unprecedented assessment of the genetic status of WBP from western central Africa. Combining mtDNA sequencing and microsatellites genotyping of several hundreds of pangolins, we identified range overlap between the WCA and Gab mitochondrial lineages with limited introgression in southern Cameroon, so the two lineages can still be considered separate evolutionary units. Further sampling and sequencing efforts –particularly through high-throughput sequencing approaches– will have to be conducted across the study region, notably in Gabon where the number of genetic samples was low, in order to assess more accurately the admixture pattern between WCA and Gab and geographic structuring within WCA. Ideally, sampling will have to be oriented towards forest sites in order to minimize the potential bias of inaccurate sourcing radius of the local markets herein used as reference populations (notably relative to a potentially artifactual Wahlund effect). Such refined data will be crucial to establish a genetic-based management unit delineation strategy for guiding the conservation of WBP in the study region. As a corollary, new knowledge on the dispersal and breeding behavior of WBP, which remains literally unknown, will have to be produced urgently.

Our set of markers provides a unique opportunity to trace the WBP trade at the individual and population levels across western central Africa, and should be considered a valuable resource for future forensic applications at reasonable costs. For the geographic tracing of WBP from western central Africa, and because single locus (mtDNA)-typing has recently been applied to trace the pangolin trade (Luczon et al., 2016; Mwale et al., 2016; Zhang et al., 2020), we recommend a multi-locus approach given the admixture pattern observed between WCA and Gab. Because introgression may occur between other neighboring lineages of WBP (Gossé et al., in prep.), it will be important in the future to rely on a global reference database where all the WBP lineages will have been genotyped.

According to our results and previous investigations, Yaoundé is a main hub for the pangolin trade in the sub-region, where WBP are openly sold on the stalls despite being nationally protected (Din Dipita et al., 2022; Harvey-Carroll et al., 2022). The Yaoundé market is driven by a large urban demand for bushmeat consumption (Edderai & Dame, 2006; Randolph, 2016; Saylors et al., 2021), and as such should receive urgent attention from researchers, conservationists and anti-poaching services as a crucial point of entry to mitigate the WBP trade in Cameroon and, more globally, western central Africa. A similar conclusion applies to European points of entry of WBP carcasses (Chaber et al., 2010; Heinrich et al., 2019; Heinrich et al., 2016), where CITES regulations are not efficiently implemented because of a global lack of prioritization, means and training at European borders (Din Dipita et al., 2022; Halbwax, 2020). As a general conclusion, better sensitization of range-state wildlife management services and foreign customs agencies to the regional and global trade of WBP should help prioritizing the conservation of the species.

## Supporting information

https://drive.google.com/file/d/1wyvaAwKhSxbutnsBFbN6Sy9reiIF-RiV/view?usp=share_link

https://drive.google.com/file/d/1fdfiwTmTJTYxsfIjJ_3YGxU3q--xL0QS/view?usp=share_link

https://docs.google.com/spreadsheets/d/1esEFV_txUayM21EJO2ZJBB0vqhuROdd7/edit?usp=share_link&ouid=105823212438779530523&rtpof=true&sd=true

## Acknowledgements

We are grateful to the Ministère de la Recherche Scientifique et de l’Innovation for providing the research permit (000094/MINRESI/B00/C00/C10/nye). We thank the staff of the “Plateau technique – Biologie moléculaire et microbiologie” at EDB for assistance during lab work. We thank Franklin and Jean for their help during field work. The local hunters and vendors gently provided us with the necessary tissue samples by free consent. We are grateful to V. Benoît and L. Cervantes (Institut de Recherche Criminelle de la Gendarmerie Nationale, Cergy Pontoise, France) for giving us access to the samples seized at Roissy airport (France). ADD, SA and PG received support from the Agence Nationale de la Recherche (ANR-17-CE02-0001; PANGO-GO). ADD and PG were also supported by the Fundação para a Ciência e Tecnologia (FCT IC&DT 02/SAICT/2017 - n° 032130; BUSHRISK). ADD received a grant « Séjours scientifiques de haut niveau » (No. 968962A) from the Service de Coopération et d’Action Culturelle (SCAC) of the Ambassade de France in Cameroon.

## Authors’ contributions

PG, ADM, EL, KA and ADD conceived the ideas; ADD, ADM, FN, PG, ALC, SN and BRM collected the samples; ADD, SA and PG conducted the molecular laboratory work; ADD, SA and PG analyzed the data; ADD and PG led the writing; ADM, SA, EL, MT, FN, BRM, KA, SN and ALC read and revised the drafted work. All the co-authors have read and approved the final version of the manuscript.

## Data availability

The sequence data obtained in this study have been deposited in GenBank (NCBI) under accession numbers OP524750 - OP525226 (https://www.ncbi.nlm.nih.gov/).

This work is original and has not been published elsewhere.

We declare that the authors have no known competing financial interests or personal relationships that could have appeared to influence the work reported in this paper.

