## Supplementary material for "Genetic tracing of the white-bellied pangolin’s trade in western central Africa": https://drive.google.com/file/d/1wyvaAwKhSxbutnsBFbN6Sy9reiIF-RiV/view?usp=share_link

Supplementary Table S1. Samples of white-bellied pangolins used in this study, including information on their geographic referencing, lineage / cluster assignment, tracing and genotypes [Excel File].
In red, individuals identified as cyto-nuclear hybrids or admixed individuals between the two lineages present in western central Africa.

Lineages as defined in Gaubert et al. (2016): WCA = Western Central Africa; Gab = Gabon.

Populations: GES = southern Equatorial Guinea; Mt Cam = National Park of Mt Cameroon; PNCM = National Park of Campo Ma'an

Supplementary Table S2. Genetic diversity estimates among white-bellied pangolin lineages based on cytochrome *b*.

| | **WCA** | ***WCA (Gaubert et al., 2016)*** | **Gabon** | **Dahomey Gap** | **Western Africa** | **Central Africa** | **Ghana** |
|---|---|---|---|---|---|---|---|
| Number of sequences | 552 | *152* | 13 | 14 | 12 | 14 | 4 |
| Sequence length (bp) | 381 | *402* | 381 | 402 | 402 | 402 | 402 |
| Number of Haplotypes, *h* | 67 | *31* | 8 | 5 | 6 | 7 | 3 |
| Haplotype diversity, *Hd* | 0.830 | *0.811* | 0.897 | 0.703 | 0.682 | 0.879 | 0.833 |
| Nucleotide diversity, $\pi$ | 0.010 | *0.006* | 0.019 | 0.003 | 0.002 | 0.007 | 0.003 |

Supplementary Table S3. Locus significantly affected by null alleles under the assumption of Hardy-Weinberg equilibrium (in bold, after Bonferroni correction), in white-bellied pangolins from the Foumbot population (WCA; N = 31).

Adjusted allele's frequencies of amplified allele's bases on the four correction methods (Oosterhout, Chakraborty, Brookfield 1 and Brookfield 2) of null allele estimation.

| Locus | Null Present | Oosterhout | Chakraborty | Brookfield 1 | Brookfield 2 |
|---|---|---|---|---|---|
| PT1162028 | no | 0,0369 | 0,039 | 0,0273 | 0,0273 |
| PT1753627 | no | -0,0448 | -0,0382 | -0,0346 | 0 |
| **PT796077** | **yes** | 0,1578 | 0,1924 | 0,1454 | 0,1454 |
| **PT1973508** | **yes** | 0,083 | 0,0904 | 0,0769 | 0,0769 |
| **PT839522** | **yes** | 0,2901 | 0,4504 | 0,2469 | 0,2469 |
| PT464918 | no | -0,0262 | -0,0208 | -0,0181 | 0 |
| **PT1453906** | **yes** | 0,1384 | 0,1748 | 0,1153 | 0,1153 |
| PT34432 | no | -0,0473 | -0,0406 | -0,0338 | 0 |
| **PT1594892** | **yes** | 0,0894 | 0,0984 | 0,0803 | 0,0803 |
| PT308752 | no | -0,0151 | -0,0134 | -0,0121 | 0 |
| PT1669238 | no | 0,0407 | 0,0485 | 0,0391 | 0,0391 |
| PT1225378 | no | -0,0203 | -0,0255 | -0,0126 | 0 |
| PT739516 | no | 0,0434 | 0,0649 | 0,0315 | 0,0315 |
| PT619913 | no | 0,0905 | 0,1021 | 0,0654 | 0,0654 |
| **PT338821** | **yes** | 0,1182 | 0,1296 | 0,0962 | 0,0962 |
| **PT1849728** | **yes** | 0,1299 | 0,172 | 0,1011 | 0,1794 |
| PT378852 | no | 0,0966 | 0,1065 | 0,0607 | 0,1553 |
| **PT353755** | **yes** | 0,2079 | 0,2678 | 0,1799 | 0,2329 |
| PT276641 | no | -0,043 | -0,027 | -0,0209 | 0 |
| PT2019332 | no | -0,0058 | -0,0007 | -0,0007 | 0 |

Supplementary Table S4. Deviation of genotypic frequencies from those expected under Hardy-Weinberg equilibrium in white-bellied pangolins from the Foumbot population.

Significant values are in bold (after Bonferroni correction).

| **Locus** | PT1162028 | PT1753627 | PT796077 | PT1973508 | PT839522 | PT464918 | PT1453906 | PT34432 | PT1594892 | PT308752 |
|---|---|---|---|---|---|---|---|---|---|---|
| | 0.667 | 0.997 | 0.055 | 0.402 | **0.000*** ** | **0.000*** ** | 0.189 | 1.000 | **0.001*** | 0.604 |

[redacted]

| **Locus** | PT1669238 | PT1225378 | PT739516 | PT619913 | PT338821 | PT1849728 | PT378852 | PT353755 | PT276641 | PT2019332 |
|---|---|---|---|---|---|---|---|---|---|---|
| | 0.091 | 0.908 | 0.015 | 0.063 | 0.188 | 0.138 | 0.347 | **0.000*** ** | 0.904 | 0.986 |

Supplementary Table S5. Pairwise differentiation estimates ($F_{ST}$) among populations of white-bellied pangolins from Cameroon, Equatorial Guinea and Gabon (significant values in bold) based on 20 microsatellites loci.

Populations refer to Table S1. Significant values are in bold.

| **Populations** | **Bayomen** | **PNCM** | **Abong Mbang** | **Eseka** | **Yabassi** | **Foumbot** | **GES** | **Manengole** | **Sangmelima** | **Mt Cam** | **Gabon** |
|---|---|---|---|---|---|---|---|---|---|---|---|
| **Bayomen** | 0.00000 | | | | | | | | | | |
| **PNCM** | **0.03976** | 0.00000 | | | | | | | | | |
| **Abong Mbang** | 0.02465 | 0.00792 | 0.00000 | | | | | | | | |
| **Eseka** | 0.00525 | 0.01817 | 0.00593 | 0.00000 | | | | | | | |
| **Yabassi** | 0.01378 | **0.04401** | **0.03349** | **0.02374** | 0.00000 | | | | | | |
| **Foumbot** | 0.00541 | **0.03251** | **0.02601** | **0.02152** | 0.01287 | 0.00000 | | | | | |
| **GES** | **0.04852** | **0.03012** | **0.03201** | **0.03550** | **0.03378** | **0.03096** | 0.00000 | | | | |
| **Manengole** | 0.01116 | **0.02952** | **0.01930** | **0.01475** | 0.01659 | **0.02317** | **0.03240** | 0.00000 | | | |
| **Sangmelima** | 0.02283 | 0.00608 | 0.00715 | 0.00862 | **0.02883** | **0.02175** | **0.02574** | **0.01550** | 0.00000 | | |
| **Mt Cam** | -0.01168 | 0.01783 | 0.01957 | 0.00610 | 0.00857 | 0.01539 | 0.01918 | -0.00132 | 0.01162 | 0.00000 | |
| **Gabon** | **0.21310** | **0.20057** | **0.18398** | **0.19074** | **0.21153** | **0.22489** | **0.19854** | **0.20439** | **0.19015** | **0.18971** | 0.00000 |

Supplementary Table S6. Private alleles from seven loci potentially usable to trace the trade of white-bellied pangolins in western central Africa. Number of individuals = 0 refers to private alleles characterizing a reference population but not found in any white-bellied pangolins from urban markets and seizures.

| **Locus** | **Population** | **Private allele (size)** | **Number of individuals assigned from urban markets and seizures** |
|---|---|---|---|
| PT1669238 | Mt Cameroon | **186** | 9 |
| PT2019332 | Abong Mbang | **186** | 2 |
| PT308752 | Manengole | **148** | 2 |
| PT308752 | Abong Mbang | **176** | 3 |
| PT308752 | South of Equatorial Guinea | **180** | 11 |
| PT308752 | Yabassi | **208** | 0 |
| PT338821 | Sangmelima | **256** | 5 |
| PT338821 | Abong Mbang | **268** | 1 |
| PT338821 | Sangmelima | **274** | 1 |
| PT338821 | Sangmelima | **282** | 1 |
| PT338821 | Sangmelima | **284** | 0 |
| PT34432 | Sangmelima | **109** | 0 |
| PT34432 | Eseka | **115** | 5 |
| PT34432 | Yabassi | **125** | 2 |
| PT34432 | Abong Mbang | **129** | 3 |
| PT34432 | Mt Cameroon | **184** | 0 |
| PT796077 | Mt Cameroon | **161** | 0 |
| PT839522 | Abong Mbang | **205** | 2 |

Supplementary Table S7. List of the 47 white-bellied pangolins from urban markets or international seizures traced to their reference populations in western central Africa.

| Locus | Assigned individual | Market / Seizure | Source population |
|---|---|---|---|
| **PT839522** | Dla A1 | Douala Central Market | Abong Mbang |
| **PT34432** | Dla B45 | Douala Dakat Market | Abong Mbang |
| **PT308752** | Dla B59 | Douala Dakat Market | Abong Mbang |
| **PT338821** | LL180926022B | Brussels Airport (Belgium) | Abong Mbang |
| **PT839522** | Y137 | Yaoundé Market | Abong Mbang |
| **PT34432** | Y146 | Yaoundé Market | Abong Mbang |
| **PT2019332** | Y216 | Yaoundé Market | Abong Mbang |
| **PT308752** | Y354 | Yaoundé Market | Abong Mbang |
| **PT308752** | Y69 | Yaoundé Market | Abong Mbang |
| **PT34432** | Y69 | Yaoundé Market | Abong Mbang |
| **PT2019332** | Y90 | Yaoundé Market | Abong Mbang |
| **PT308752** | Y100 | Yaoundé Market | Manengole |
| **PT308752** | Y359 | Yaoundé Market | Manengole |
| **PT1669238** | Dla B13 | Douala Dakat Market | Mt Cameroon |
| **PT1669238** | Dla B85 | Douala Dakat Market | Mt Cameroon |
| **PT1669238** | Y109 | Yaoundé Market | Mt Cameroon |
| **PT1669238** | Y179 | Yaoundé Market | Mt Cameroon |
| **PT1669238** | Y251 | Yaoundé Market | Mt Cameroon |
| **PT1669238** | Y33 | Yaoundé Market | Mt Cameroon |
| **PT1669238** | Y349 | Yaoundé Market | Mt Cameroon |
| **PT1669238** | Y361 | Yaoundé Market | Mt Cameroon |
| **PT1669238** | Y64 | Yaoundé Market | Mt Cameroon |
| **PT338821** | Y129 | Yaoundé Market | Sangmelima |
| **PT338821** | Y352 | Yaoundé Market | Sangmelima |
| **PT338821** | Y360 | Yaoundé Market | Sangmelima |
| **PT338821** | Y374 | Yaoundé Market | Sangmelima |
| **PT338821** | Y77 | Yaoundé Market | Sangmelima |
| **PT338821** | Y2 | Yaoundé Market | Sangmelima |
| **PT34432** | Dla B102 | Douala Dakat Market | Sangmelima |
| **PT308752** | LL180926022A | Brussels Airport (Belgium) | South of Equatorial Guinea |
| **PT308752** | Y162 | Yaoundé Market | South of Equatorial Guinea |
| **PT308752** | Y221 | Yaoundé Market | South of Equatorial Guinea |
| **PT308752** | Y222 | Yaoundé Market | South of Equatorial Guinea |
| **PT308752** | Y228 | Yaoundé Market | South of Equatorial Guinea |
| **PT308752** | Y229 | Yaoundé Market | South of Equatorial Guinea |
| **PT308752** | Y336 | Yaoundé Market | South of Equatorial Guinea |
| **PT308752** | Y342 | Yaoundé Market | South of Equatorial Guinea |
| **PT308752** | Dla B100 | Douala Dakat Market | South of Equatorial Guinea |
| **PT308752** | Dla B101 | Douala Dakat Market | South of Equatorial Guinea |
| **PT308752** | Y332 | Yaoundé Market | South of Equatorial Guinea |
| **PT34432** | Y157 | Yaoundé Market | Eseka |
| **PT34432** | Y196 | Yaoundé Market | Eseka |
| **PT34432** | Y232 | Yaoundé Market | Eseka |
| **PT34432** | Y242 | Yaoundé Market | Eseka |

| Locus | Assigned individual | Market / Seizure | Source population |
|---|---|---|---|
| **PT34432** | Dla A30 | Douala Central Market | Yabassi |
| **PT34432** | Dla B3 | Douala Dakat Market | Yabassi |
| **PT34432** | Y96 | Yaoundé Market | Eseka |

Supplementary Figure S1. Map of the 40 bushmeat markets and forest sites surveyed in Cameroon, Equatorial Guinea and Gabon.

S1-Abong Mbang; S2-Akonolinga; S3-Bangong; S4-Bayib Assibong; S5-Bioko; S6-Bipindi; S7-Campo; S8-Douala Central Market; S9-Douala Dakat Market; S10-Djoum; S11-Ekombitié; S12-Eseka; S13-Esse; S14-Foumbot; S15-Bitam; S16-Anguma; S17-Bongoro; S18-Ebenguan; S19-Emangos; S20-Misergue; S21-Taguete; S22-Makokou; S23-Medouneu; S24-Lolodorf; S25-Maan; S26-Mamb; S27-Manengole; S28-Manyemen; S29-Nditam; S30-Sangmelima; S31-Seizure; S32-Yabassi; S33-Yaoundé Nkolndongo Market; S34-Yokadouma; S35-Franceville; S36-Mocabe; S37-Nkoltang; S38-Okoumbi; S39-Oyem; S40-Tchibanga.

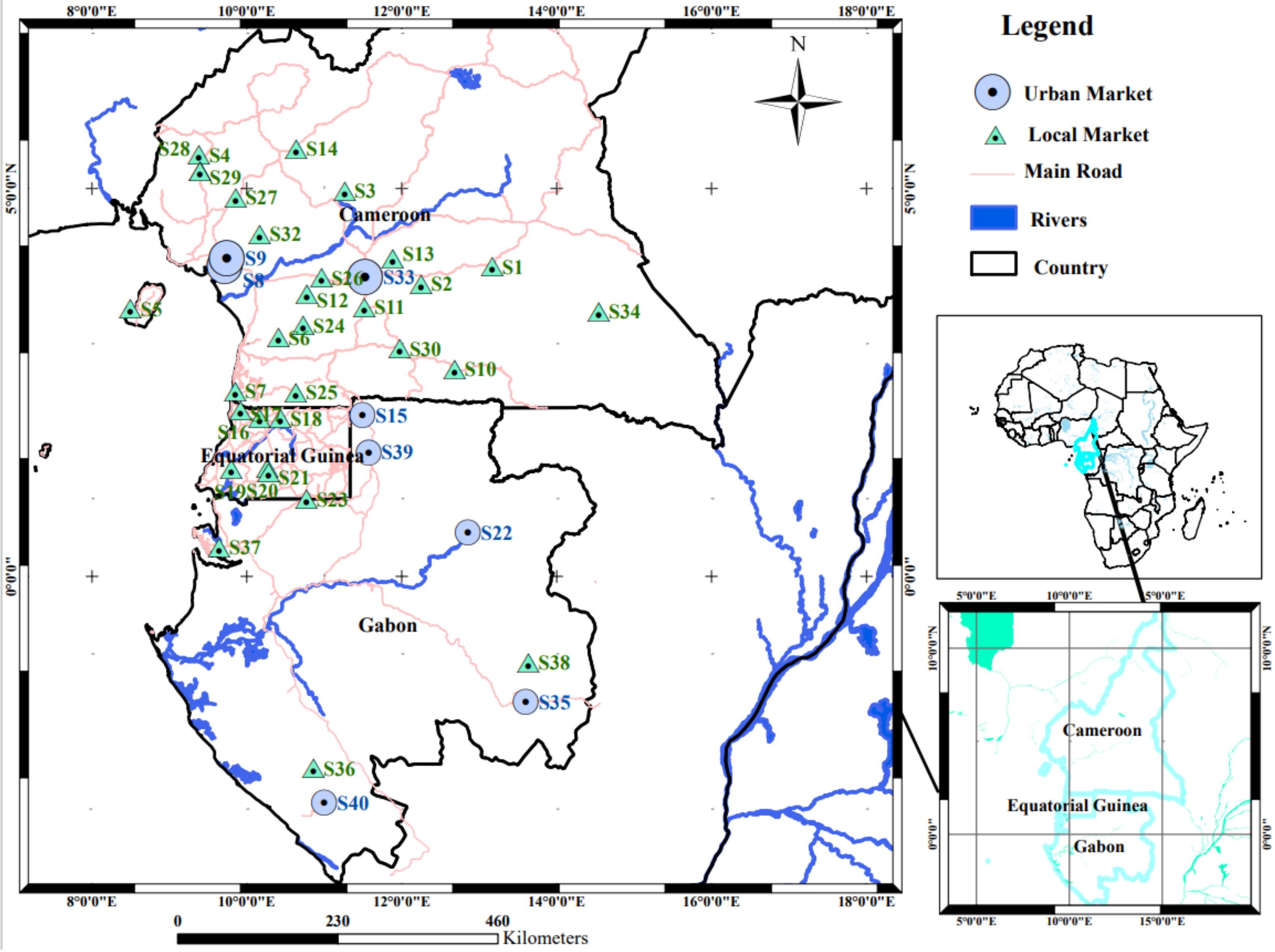
8°0'0"E
10°0'0"E
12°0'0"E
14°0'0"E
16°0'0"E
18°0'0"E
5°0'0"N
0°0'0"
Legend
Urban Market
Local Market
Main Road
Rivers
Country
N
Cameroon
Equatorial Guinea
Gabon
S1
S2
S3
S4
S5
S6
S7
S8
S9
S10
S11
S12
S13
S14
S15
S16
S17
S18
S19
S20
S21
S22
S23
S24
S25
S26
S27
S28
S29
S30
S32
S33
S34
S35
S36
S37
S38
S39
S40
0
230
460
Kilometers
5°0'0"E
10°0'0"E
15°0'0"E
10°0'0"N

Supplementary Figure S2. Neighbor joining tree of white-bellied pangolins based on 626 cytochrome *b* sequences. [pdf file]

Bootstrap values > 75% are shown at nodes. Sequence clusters are taxonomically delimitated according to Gaubert et al. (2016). Sample numbering corresponds to Table S1. Scale bar at bottom represents percentage of K2P distance. **** designates the outgroup.

Supplementary Figure S3. Median-joining network showing mutational relationships between the 74 haplotypes observed in 553 white-bellied pangolins from Cameroon (yellow), Equatorial Guinea (continent; blue), Bioko Island (green), Roissy airport – France (grey), Brussels airport – Belgium (purple), and Gabon (pink).

Size of circle is proportional to the number of haplotypes (e.g., H_30 = 1). Mutational steps are illustrated as dashes perpendicular to network connections.

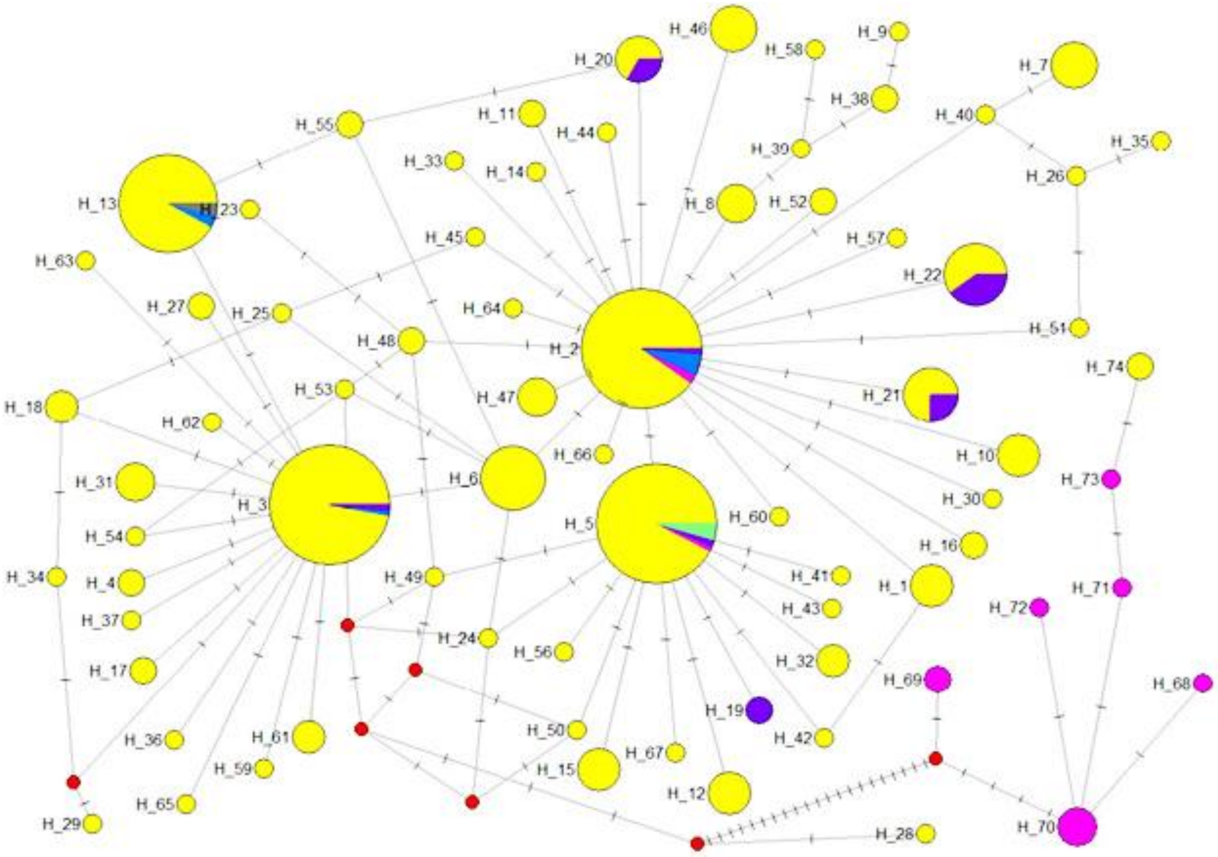

Supplementary Figure S4. Distribution of cytochrome *b* haplotypes among white-bellied pangolins from the Western Central Africa and Gabon lineages.

Site numbers refer to Figure S1. H1-H67 = Western Central Africa; H68-H74 = Gabon.

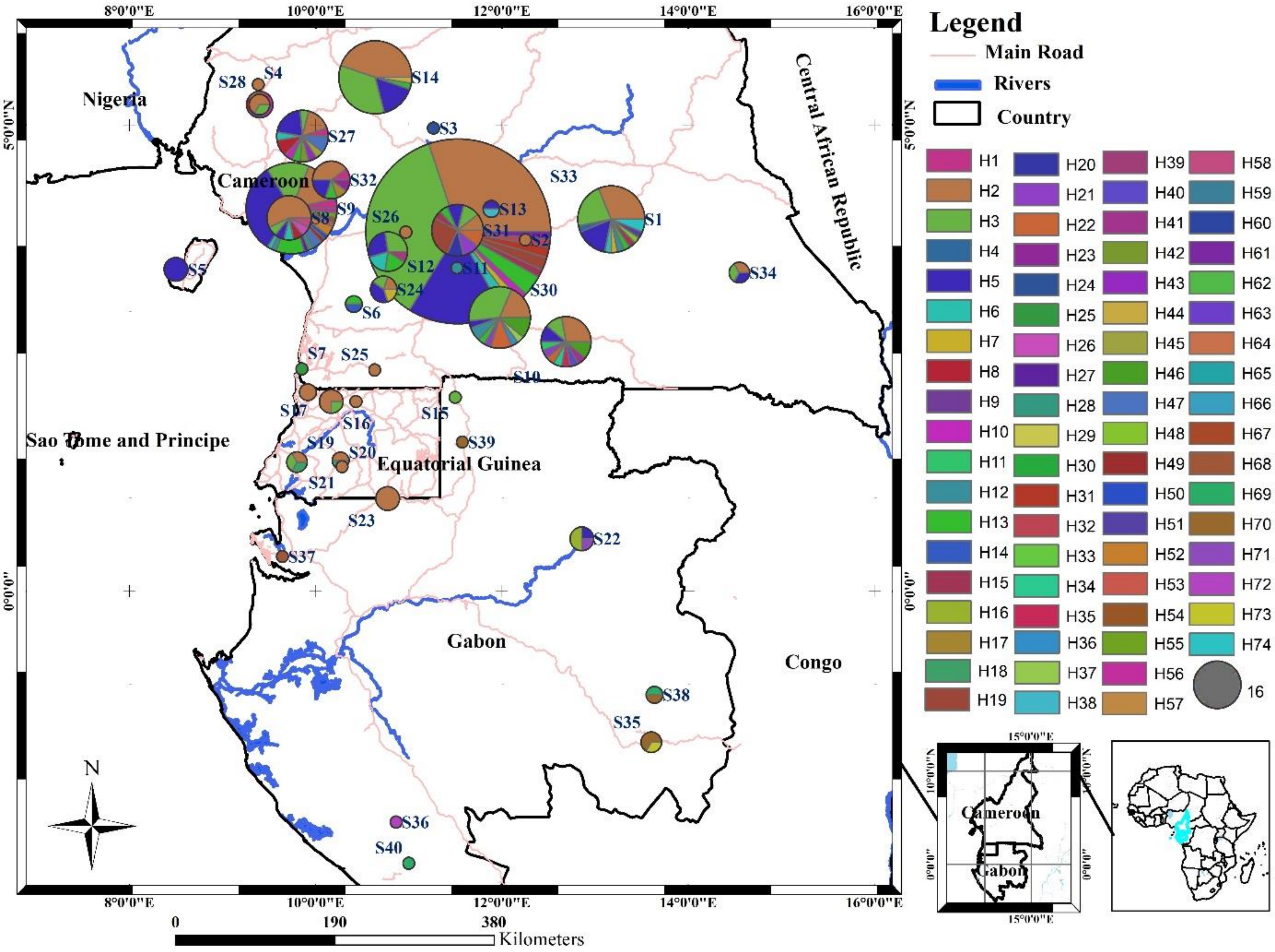

Legend
Main Road
Rivers
Country
Nigeria
Cameroon
Central African Republic
Sao Tome and Principe
Equatorial Guinea
Gabon
Congo
8°0'0"E
10°0'0"E
12°0'0"E
14°0'0"E
16°0'0"E
5°0'0"N
0°0'0"
0
190
380
Kilometers
N
16

Supplementary Figure S5. Plot of *mismatch distribution* for white-bellied pangolins from WCA under hypothesis of sudden demographic expansion (left) and spatial demographic expansion (right).

Frequencies expected in red; Frequencies observed in green.

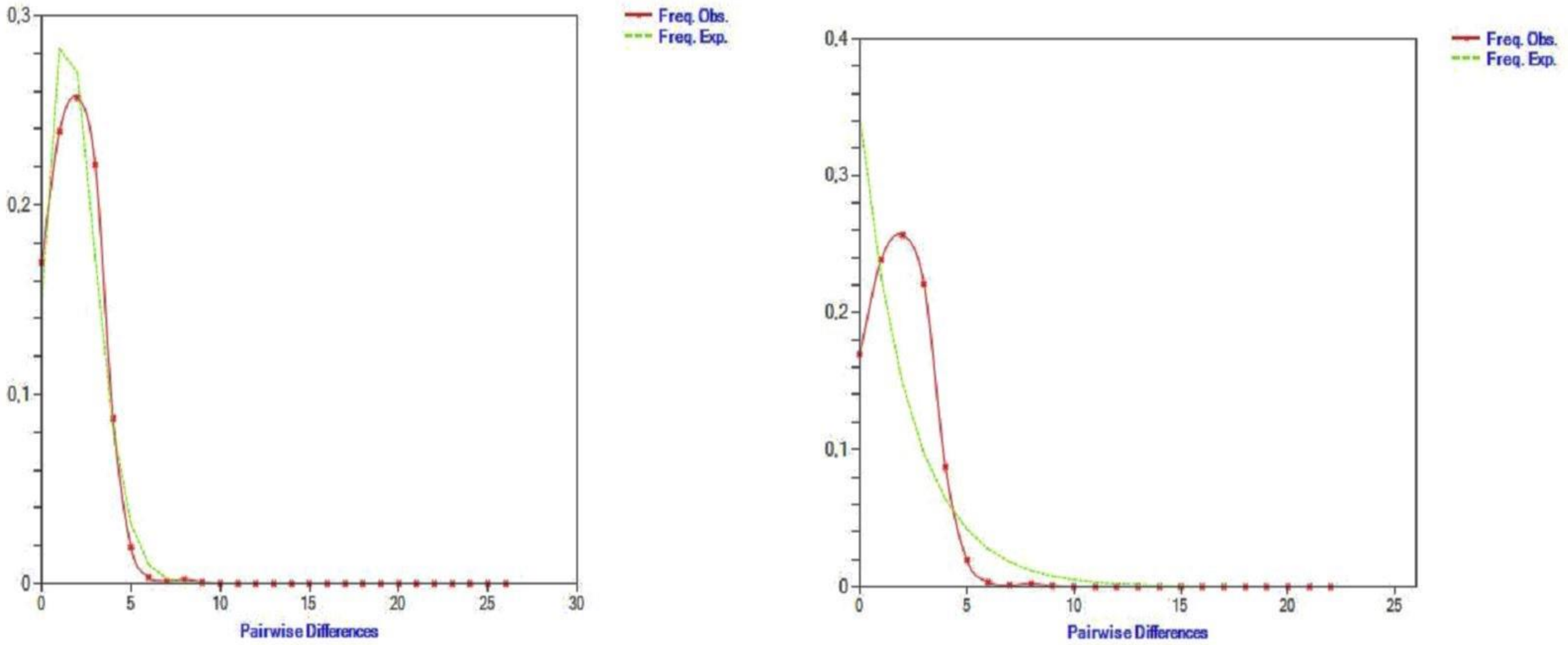

Supplementary Figure S6. Distribution of genetic variance (PCoA) among populations of white-bellied pangolins from WCA and Gab lineages.

(**a**) = including urban markets; (**b**) = excluding urban markets. Percentage of explained variance per axis is given between brackets.

Population names refer to Table S1. DM = Douala Markets; YNM = Yaoundé Market.

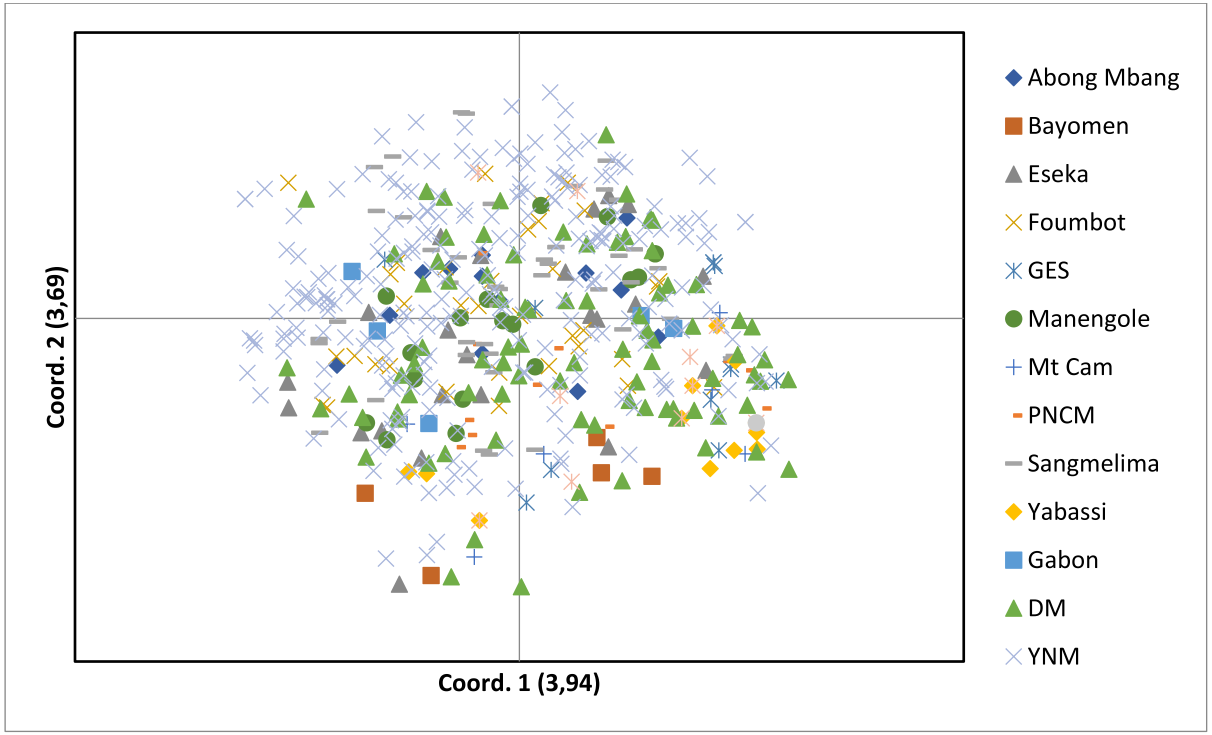


(**a**)

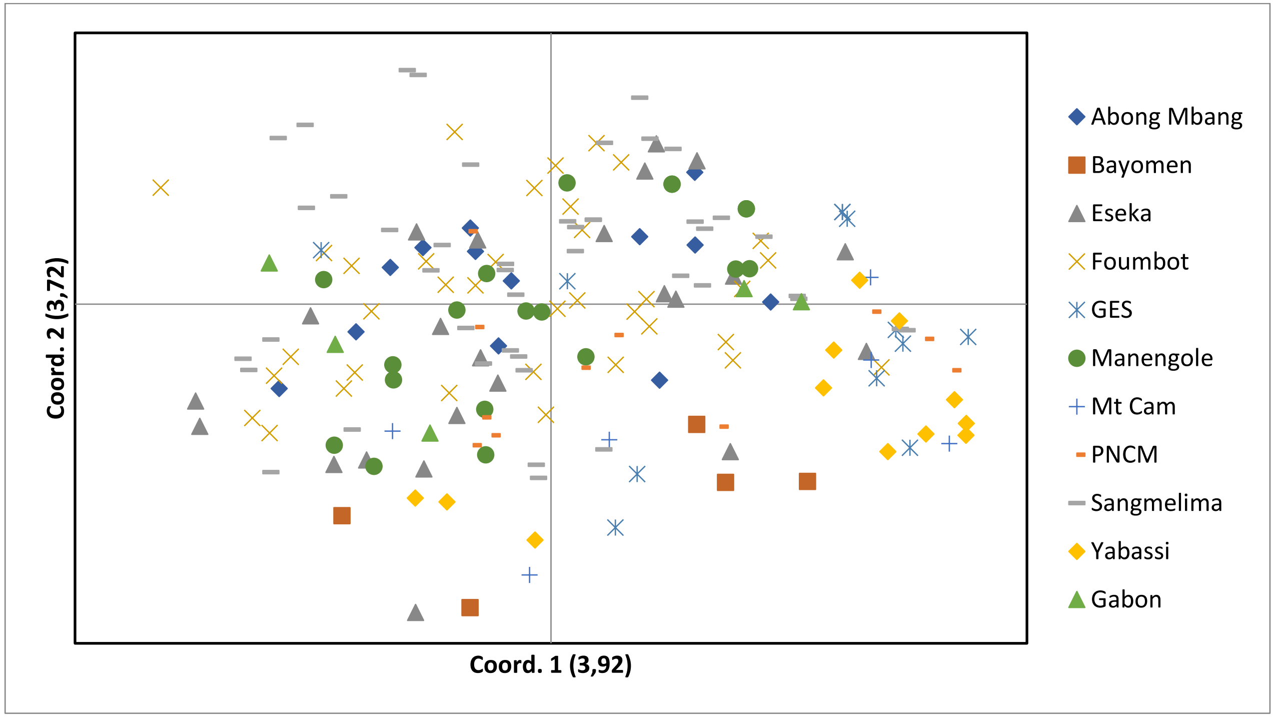
Coord. 2 (3,72)
Coord. 1 (3,92)
Abong Mbang
Bayomen
Eseka
Foumbot
GES
Manengole
Mt Cam
PNCM
Sangmelima
Yabassi
Gabon

**(b)**

Supplementary Figure S7. Assignment plots among white-bellied pangolins from western central Africa (N = 558) as assessed with STRUCTURE for K = 2 to K = 10. All the samples from urban markets, seizures and reference populations were included.

Each individual is represented by a vertical bar. The colour of STRUCTURE plot represents nuclear clusters concordant with mitochondrial lineage assignment (blue = WCA; orange = Gab). 1- Individuals mtDNA-assigned to WCA lineage; 2- Individuals mtDNA-assigned to Gab lineage.

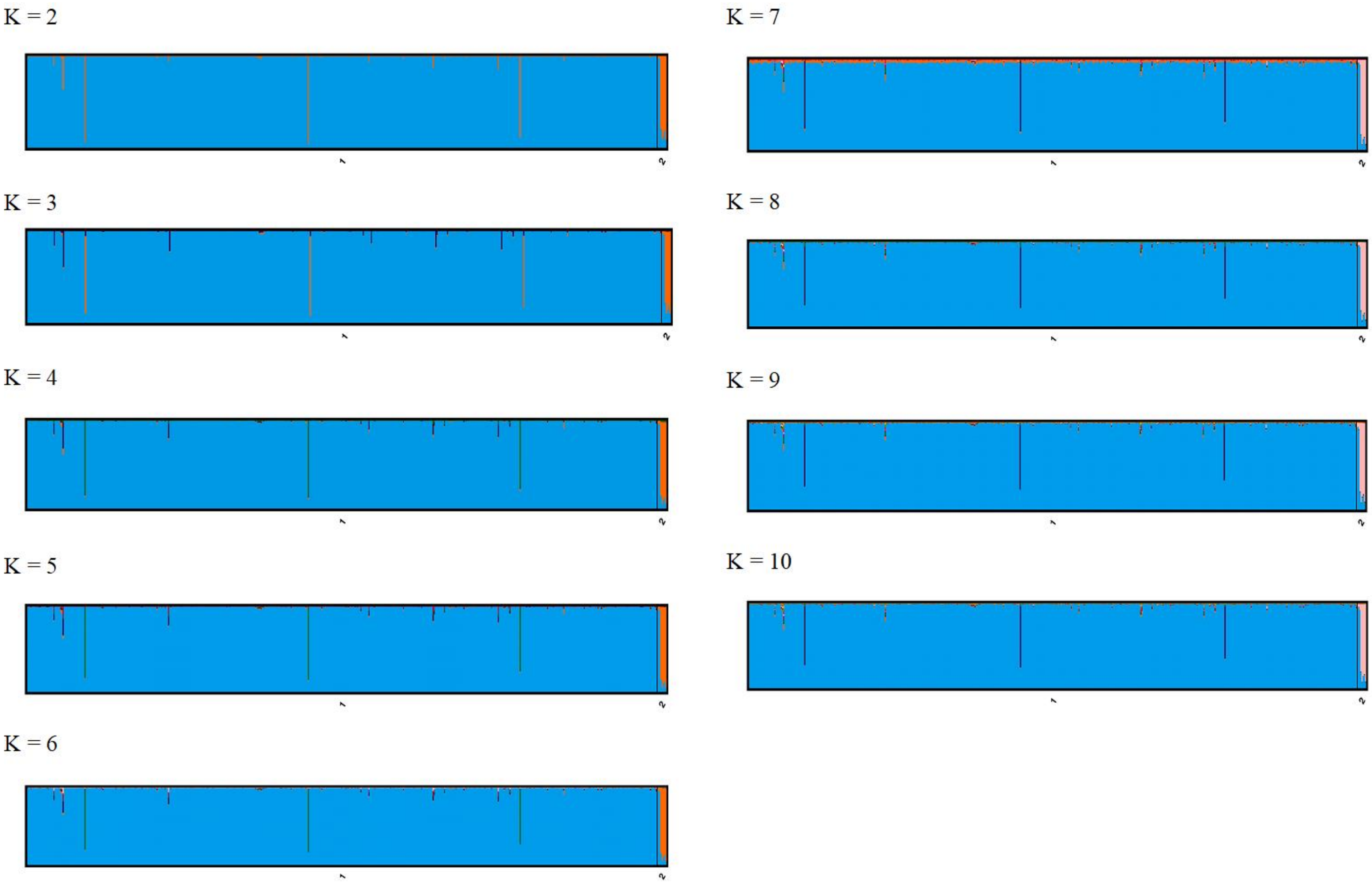
K = 2
K = 3
K = 4
K = 5
K = 6
K = 7
K = 8
K = 9
K = 10

Supplementary Figure S8. Assignment plots among white-bellied pangolins from the WCA lineage (N = 181) as assessed with STRUCTURE for K = 2 to K = 10. Only samples from 10 reference populations were included.

Each individual is represented by a vertical bar. Reference populations: 1-Bayomen; 2-PNCM; 3-Abong Mbang; 4-Eseka; 5-Yabassi; 6-Foumbot; 7-GES; 8-Manengole; 9-Sangmelima; 10-Mt Cam.

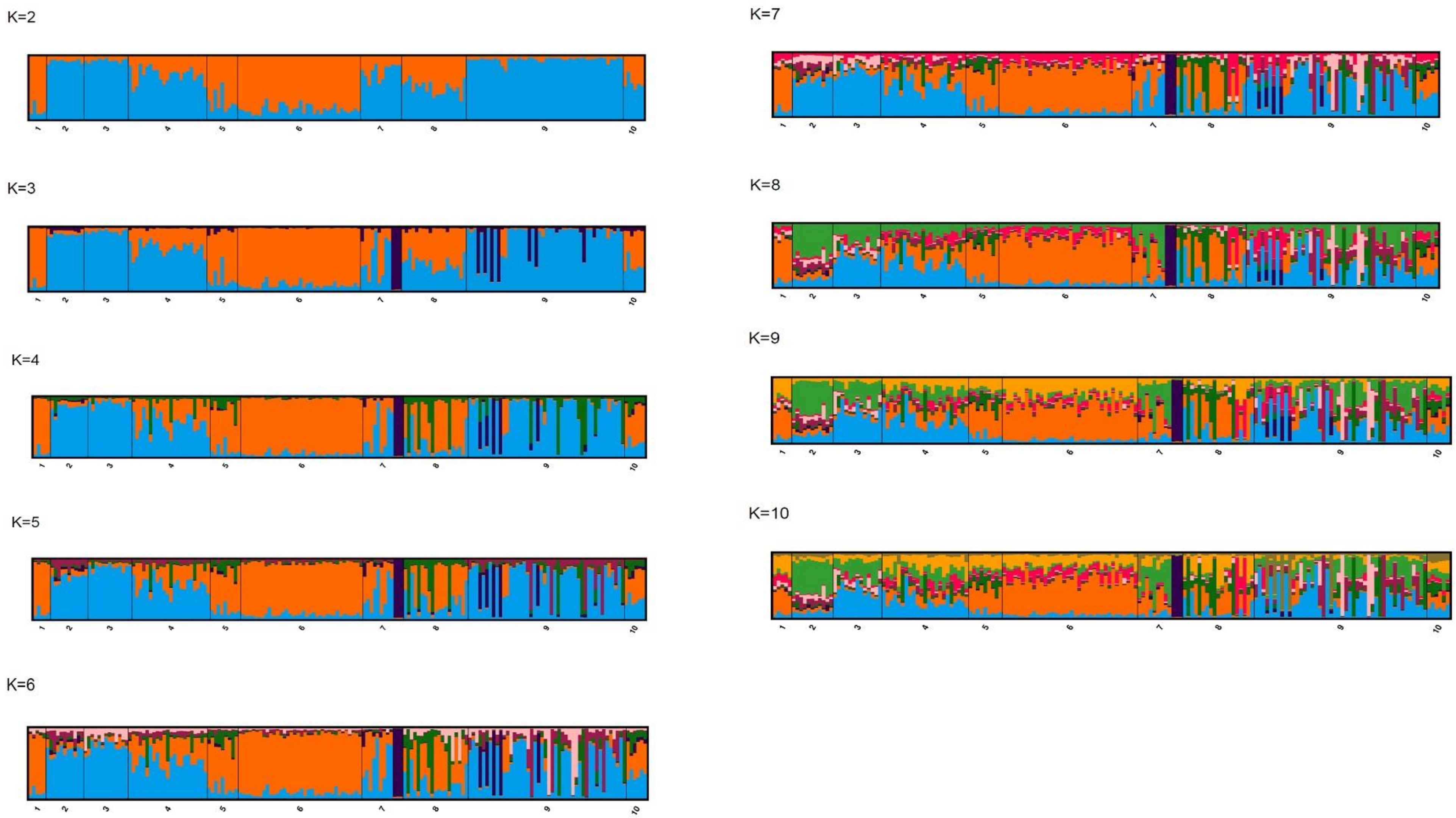
K=2
1 2 3 4 5 6 7 8 9 10
K=3
1 2 3 4 5 6 7 8 9 10
K=4
1 2 3 4 5 6 7 8 9 10
K=5
1 2 3 4 5 6 7 8 9 10
K=6
1 2 3 4 5 6 7 8 9 10
K=7
1 2 3 4 5 6 7 8 9 10
K=8
1 2 3 4 5 6 7 8 9 10
K=9
1 2 3 4 5 6 7 8 9 10
K=10
1 2 3 4 5 6 7 8 9 10

Supplementary Figure S9. Spatial clustering of white-belled pangolins in the WCA lineage obtained using *Geneland* package in R, for K = 11. Coloured areas refer to the geographic or spatial spread of the individuals; Black dots represent individuals

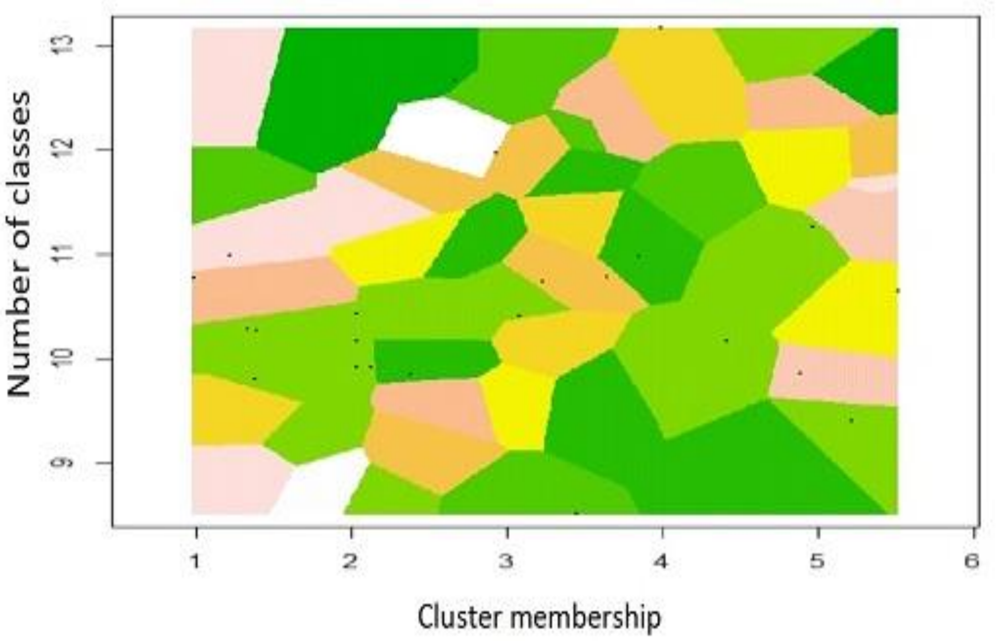

Supplementary Figure S10. Isolation by distance among (**a**) individuals and (**b**) populations of white-bellied pangolins from continental Western Central Africa as inferred from 20 microsatellites loci.

Dashed curve indicates linear regression. Dgen-Genetic distance; Dgeo-Geographic distance.

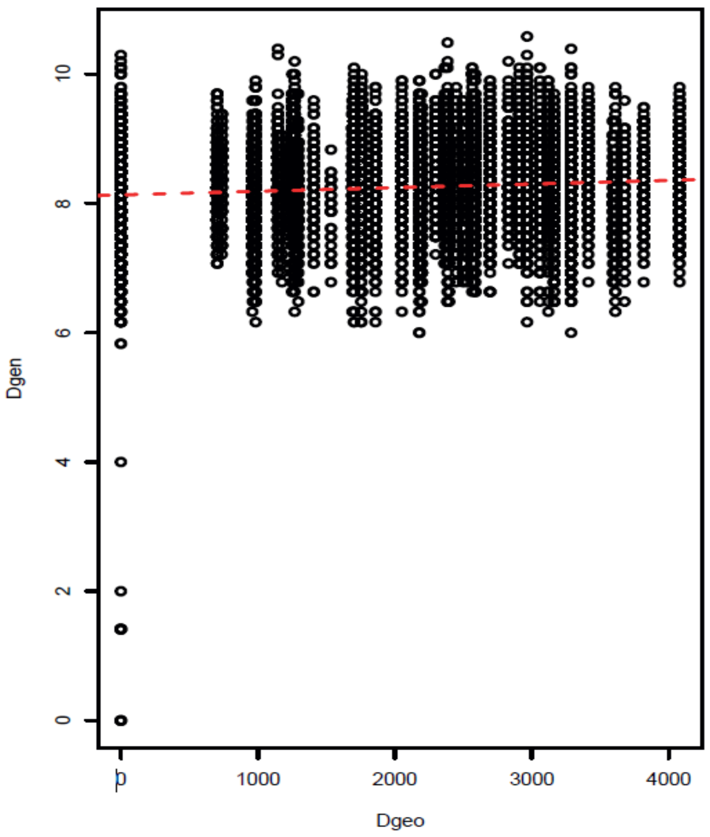


**a**

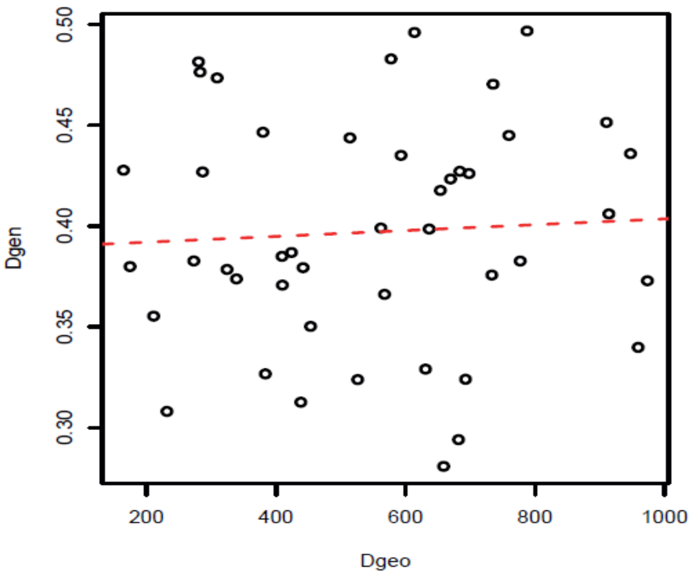


**b**

Supplementary Figure S11. Unbiased probability of identity (uPI) and probability of identity among siblings (PIsibs) for increasing, optimized combinations among the 20 microsatellite markers.

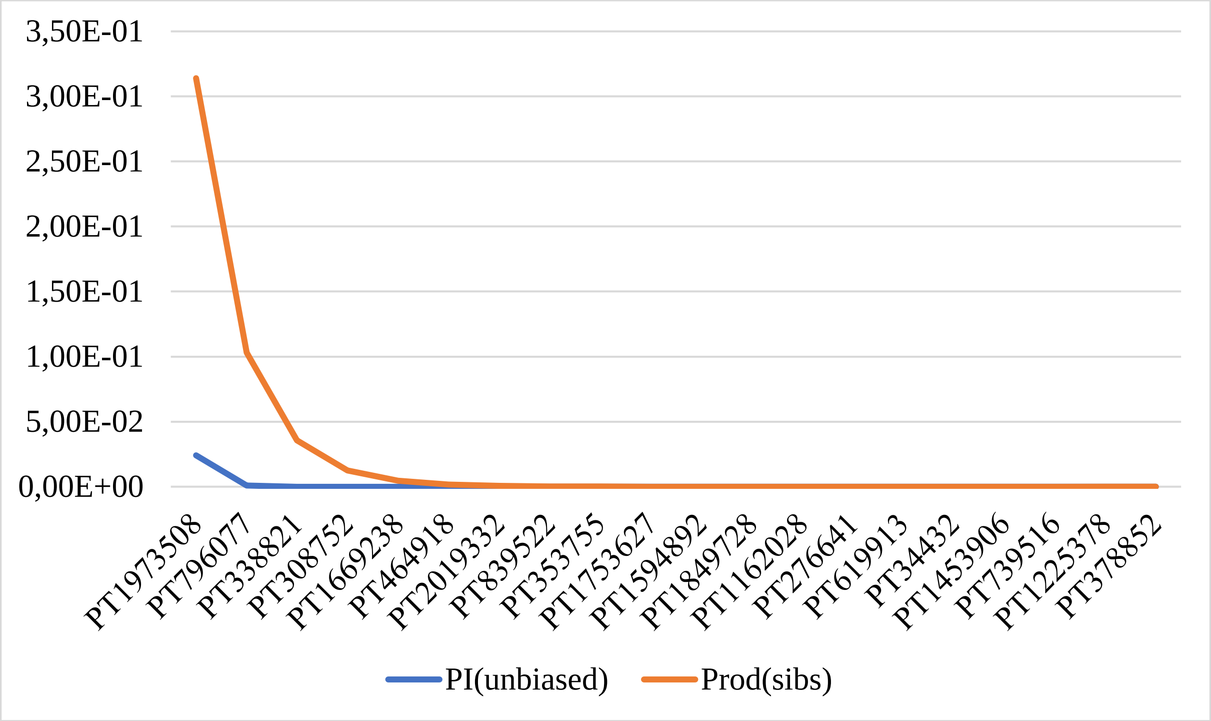
