## Supplementary figures and images for "Genetic tracing of the white-bellied pangolin’s trade in western central Africa"

### https://drive.google.com/file/d/1fdfiwTmTJTYxsfIjJ_3YGxU3q--xL0QS/view?usp=share_link

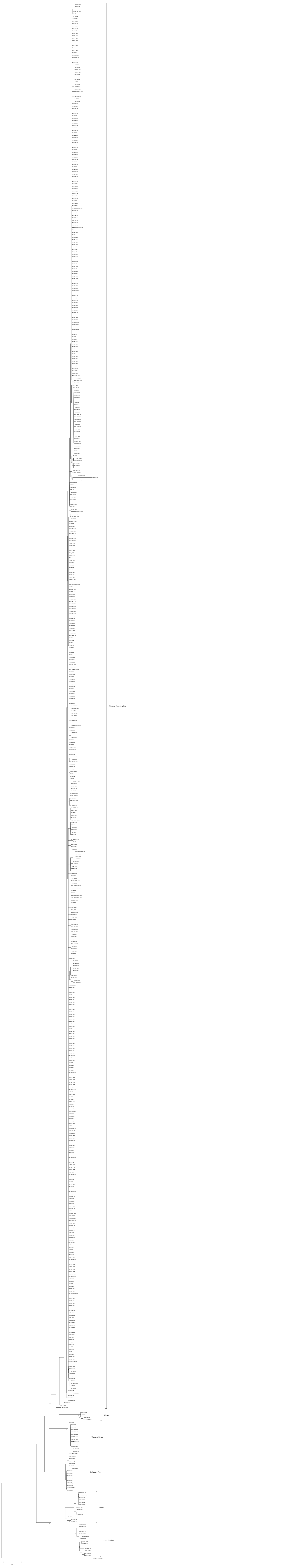
